# Mutations in Fibronectin Dysregulate Chondrogenesis in Skeletal Dysplasia

**DOI:** 10.1101/2023.12.22.573039

**Authors:** Neha E. H. Dinesh, Justine Rousseau, Deane F. Mosher, Mike Strauss, Jeannie Mui, Philippe M. Campeau, Dieter P. Reinhardt

## Abstract

Fibronectin (FN) is an extracellular matrix glycoprotein essential for the development and function of major vertebrate organ systems. Mutations in FN result in an autosomal dominant skeletal dysplasia termed corner fracture-type spondylometaphyseal dysplasia (SMDCF). The precise pathomechanisms through which mutant FN induces impaired skeletal development remain elusive. Here, we have generated patient-derived induced pluripotent stem cells as a cell culture model for SMDCF to investigate the consequences of FN mutations on mesenchymal stem cells (MSCs) and their differentiation into cartilage-producing chondrocytes. In line with our previous data, FN mutations disrupted protein secretion from MSCs, causing a notable increase in intracellular FN and a significant decrease in extracellular FN levels. Analyses of plasma samples from SMDCF patients also showed reduced FN in circulation. FN and endoplasmic reticulum (ER) protein folding chaperones (BIP, HSP47) accumulated in MSCs within ribosome-covered cytosolic vesicles that emerged from the ER and transitioned into lysosomes. Massive amounts of these vesicles were not cleared from the cytosol. The accumulation of intracellular FN and ER proteins elevated cellular stress markers and altered mitochondrial structure. Bulk RNA sequencing revealed a specific transcriptomic dysregulation of the patient-derived cells relative to controls. Analysis of MSC differentiation into chondrocytes showed impaired mesenchymal condensation, reduced chondrogenic markers, and compromised cell proliferation in mutant cells. FN mutant cells also displayed altered FN splice variants under chondrogenic stimuli. Moreover, FN mutant cells exhibited significantly lower transforming growth factor beta-1 (TGFβ1) expression, crucial for mesenchymal condensation. Exogenous FN or TGFβ1 supplementation effectively improved the MSC condensation and promoted chondrogenesis in FN mutant cells. These findings demonstrate the cellular consequences of FN mutations in SMDCF and explain the molecular pathways involved in the associated altered chondrogenesis.

**Significance /Highlights:** • SMDCF-causing mutations in fibronectin impair protein secretion in iPSC-derived mesenchymal stem cells.

• Mutant fibronectin and ER protein folding chaperones are directly exported from the rough endoplasmic reticulum into vesicles covered with ribosomes, which transition into lysosomes.

• The cells cannot clear the massive accumulation of cytosolic vesicles.

• Mutations in fibronectin impair stem cell proliferation, mesenchymal condensation, and the differentiation of MSCs into chondrocytes.

• Exogenous addition of purified fibronectin or TGFβ-1 improves mesenchymal condensation and chondrogenesis of the FN mutant stem cells.

## Introduction

Fibronectin (FN) is a master extracellular matrix organizer that mediates pivotal roles in regulating cellular and matrix-associated processes essential for the development of vertebrates [1–5]. In humans, FN is encoded by the 75 kb fibronectin gene *FN1* located on chromosome 2 [6], which gives rise to the ∼8kb transcribed premature FN mRNA. This mRNA undergoes extensive alternative splicing of exons coding for three type III domains, the extra domains A and B (EDA and EDB), and the variable domain (V), to produce the 27 known transcripts [7]. The various FN isoforms are critical determinants of physiological and pathological processes associated with major organ systems, such as embryonic development, neuronal patterning, development and maintenance of lungs, mammary glands, blood vasculature, fibrosis, cancer, and others [8–11]. FN is secreted from cells as a 440 kDa dimer linked via intermolecular disulfide bonds at the C-terminal end. Structurally, each of the 220 kDa FN monomers is composed of tandem arrays of three distinct types of domains: the type I domains (FNI1–12), type II domains (FNII1–2), and type III domains (FNIII1–15) [12]. Global deletion in the *Fn1*^-/-^ mouse model results in embryonic lethality with severe defects in developmental patterning, presence of neural tube kinks, abnormal mesoderm development, reduced embryo size, and severe growth retardation [13]. Another mutant mouse model with inactivation of the FN RGD integrin-binding motif (FN^RGE/RGE^) displayed overlapping defects [14]. Additionally, these mice displayed shortening of the posterior trunk and absence of limb buds during early embryogenesis [14]. Mutations in FN were first reported in 2008, leading to glomerulopathy with fibronectin deposits (GFND) (OMIM #601894) [15]. We and others reported heterozygous autosomal dominant mutations in *FN1* that cause a rare form of skeletal dysplasia termed “corner fracture” type spondylometaphyseal dysplasia (SMDCF) (OMIM #184255) [7,16–18]. Affected SMDCF patients suffer from visible clinical implications such as short stature and overall growth defects, severe scoliosis, coxa vara, abnormal ossification of the metaphyses, and the presence of vertebral anomalies [16]. Notably, these FN mutations only affect the skeletal system in SMDCF patients but no other organ systems.

Skeletal development in humans begins during embryogenesis when undifferentiated MSCs condense to form the cartilage anlagen, which eventually remodel during embryonic growth to give rise to the appendicular skeleton by endochondral ossification [19]. This is a highly orchestrated process, and perturbations in endochondral ossification due to genetic mutations lead to skeletal abnormalities and dysplasias [19,20]. FN and several other ECM proteins are critical components of the developing growth plate and regulate mesenchymal condensation and endochondral ossification [19]. FN is highly expressed in mesenchymal limb buds and during MSC condensation [21]. A previous study reported that assembled FN promoted MSCs to condense and a transient knockdown of FN in MSCs impaired cell condensation [22]. Additionally, FN promoted chondrogenesis in several other *in vitro* studies and is also implicated in having a role in cartilage-associated pathologies (for review, see [7]). However, its exact functional role during MSC-to-chondrocyte differentiation, skeletal development, and growth, as well as in the pathomechanisms underlying SMDCF, remains poorly understood.

The study aims to investigate the molecular mechanisms underlying FN mutations that cause SMDCF and how impaired FN affects the functionality and differentiation of stem cells. We have employed SMDCF patient-based induced pluripotent stem cells (iPSCs) and iPSC-derived MSCs and chondrocytes. We have identified that mutations in FN cause a spectrum of cellular and matrix-associated defects in FN mutant cells that impair the mesenchymal condensation of stem cells, leading to dysregulated chondrogenesis. We have also identified the prechondrogenic regulator TGFβ1 as an essential player in this mechanism.

## Results

### Generation of a cell culture SMDCF model system

To investigate the molecular consequences of heterozygous FN mutations on MSC function and differentiation, we generated a cell culture SMDCF model system. Skin fibroblasts obtained from skin punch biopsies of two SMDCF patients with *FN1* mutations c.367T>C (FN*_C123R_*) and c.693C>G (FN*_C231W_*) and from an unaffected individual (donor details in **Supp.** Figure 1A) were used to generate iPSCs. These cells were further differentiated into MSCs and chondrocytes depending on the experimental setup (**Figure 1A**). In a baseline characterization, over 90% of cells for all iPSC groups showed robust expression of the pluripotency markers SOX2, NANOG, C-MYC, and OCT4 (**Figure 1B**). Sanger sequencing was performed on amplified cDNA samples of the patient and control MSCs to confirm the FN mutations. FN*_C123R_*mutant cells showed a heterozygous peak at nucleotide position c.367T>C confirming the FN mutation p.Cys123Arg, and FN*_C231W_* cells showed heterozygous peaks at nucleotide position c.693C>G confirming the FN mutation p.Cys231Trp in these cells (**Figure 1C**). After differentiation of the iPSCs into MSCs, over 95% of the differentiated cells of all cell groups were positive for CD73 and CD105 and were entirely negative for the undifferentiated cell marker TRA-1-60 (**Figure 1D**).

**Figure 1.**
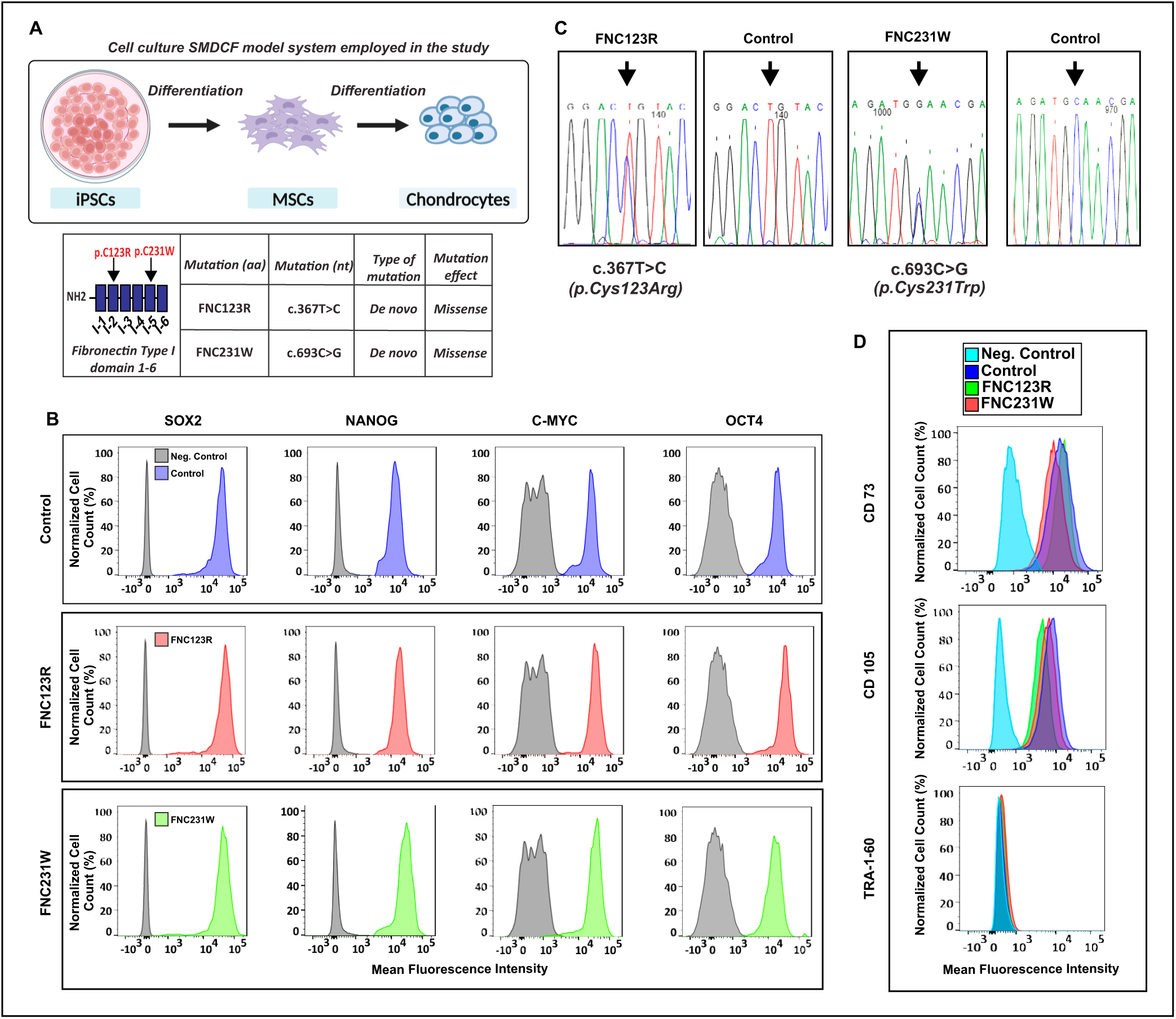
Generation of an SMDCF cell model system. **(A)** Schematic overview of an SMDCF model system using iPSCs with SMDCF-causing FN mutations. iPSCs were cultured and differentiated into MSCs and then into chondrocytes. The two FN mutations analyzed in this study are p.C123R, affecting the FNI-2 domain, and p.C231W affecting the FNI-5 domain. **(B)** iPSCs were evaluated by flow cytometry for the expression of pluripotent stem cell markers with specific antibodies against SOX2, NANOG, C-MYC, and OCT4. Positive control cell populations are represented in blue, FN*_C123R_* mutants in red and FN*_C231W_* mutants in green. Fluorescence minus one (FMO, grey) for each of the antibody was used as the negative control per cell group. **(C)** Sequencing of *FN1* mRNA from iPSC-derived MSCs after amplification by PCR, confirming the heterozygous FN mutations as indicated by arrows. **(D)** Analysis of iPSC-derived MSCs for the expression of MSC markers CD73, CD105, and undifferentiated marker TRA-1-60. Positive control cell populations are represented in blue, FN*_C123R_* mutants in green, and FN*_C231W_* mutants in red. Fluorescence minus one (FMO, cyan) for each of the antibodies was used as the negative control per cell group.

### Secretion of FN in mutant cells is impaired, leading to intracellular accumulation in vesicular structures

We previously generated recombinant mutant FN in HEK293 cells, which exhibited impaired protein secretion [16]. This prevented the structural analysis of purified mutant FN. Therefore, we now employed *in silico* structure prediction tools (Alphafold2.0, PyMOL2) to compare the known FN type I (FNI) domain structure to computationally predicted mutant FN structures for FN*_C123R_* and FN*_C231W_*. The extent of structural differences was measured in root mean square deviation (RMSD) of atomic positions of the superimposed structures. The more the RMSD values deviate from 0, the larger the structural changes. The mutant FNI-2*_C123R_* and FNI-5*_C231W_*domains showed only subtle conformational changes relative to the wild-type FNI domains, with low RMSD values of 0.73 Å and 0.82 Å, respectively (**Figure 2A**). Minor alterations were also observed for the two adjacent domains, FNI-3 (RMSD: 0.43 Å) and FNI-4 (RMSD: 0.46 Å) (**Figure 2A**). We analyzed seven additional SMDCF-causing FN mutations and obtained similar results with RMSD values ranging from 0.23-0.91 Å for the affected FNI domains and RMSD values of 0.05-0.458 Å for adjacent domains **(Supp.** Figure 1B,C). It is important to note that eight of these nine analyzed SMDCF mutations affect cysteine residues and thus disrupt intramolecular disulfide bonds. Although these missense mutations are not predicted to cause significant structural changes in the individual FNI domain structures, detrimental effects may arise from improper intra-or intermolecular disulfide bond formation that cannot be predicted using AlphaFold.

**Figure 2.**
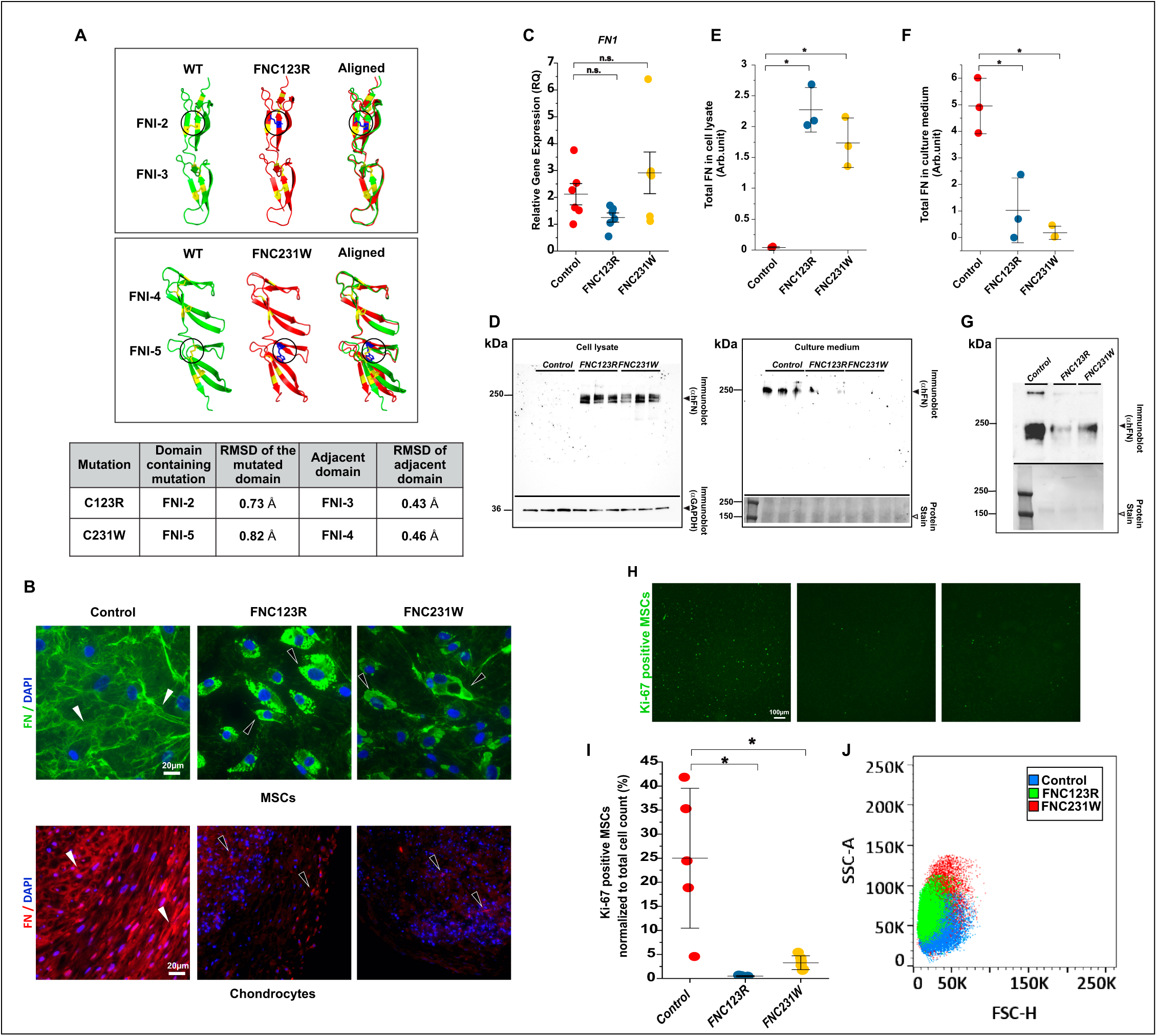
SMDCF mutations in FN cause cellular and matrix defects. **(A)** *In silico* structure prediction and comparison of the wild-type and FN mutant domains (FNI-2, FNI-3, FNI-4, FNI-5) using AlphaFold 2.0 and PyMOL. **(B)** Immunostaining images for FN in an unaffected control and the FN mutant MSCs (FN in green, top panel) and in chondrocyte micromasses (FN in red, bottom panel) using an anti-fibronectin antibody. Figure shows representative images from three independent experiments (n=3). Matrix staining for FN is denoted by black arrowheads and intracellular staining by white arrowheads. Nuclear counterstaining using DAPI is represented in blue. Relative to MSCs in monolayers, cells in micromasses show a significantly smaller cell size. **(C)** Relative gene expression of *FN1* in control and mutant MSCs. Each data point in the graph represents the average of technical triplicates or duplicates from one experiment of a total of 6 independent experiments (n=6). Error bars represent ± SEM. Statistical analysis was performed using one-way ANOVA with Bonferroni post-test. **(D)** Protein lysates and condition media from the same control and mutant MSC cultures immunoblotted for FN (∼220-250 kDa) under reducing conditions. GAPDH (37kDa) was used as the internal loading control for the cell lysate. The blot reversibly stained for proteins after the transfer was used as loading control for the culture media (unknown secreted protein, arrowheads). **(E-F)** Quantification of immunoblots shown in **(D)** from three independent experiments (n=3). **(G)** Immunoblotting for FN using serum samples obtained from an unaffected individual (*Control*) and the SMDCF patients with the FN*_C123R_* and FN*_C231W_* mutations (top panel). The blot reversibly stained for proteins was used as a loading control using an unknown visible ∼160 KDa protein (bottom panel). **(H-I)** Immunostaining of proliferation marker Ki-67 (green) using control and mutant MSCs. Ki-67 positive cells were quantified and normalized to the total cell count. Each data point represents the average of normalized Ki-67 positive cells from individual images of one representative experiment. 4 independent experiments were performed with similar results. The error bar represents ± SD. Statistical analysis was performed using one-way ANOVA with Bonferroni post-test. **(J)** Side scatter area (SSC-A) versus forward scatter height (FSC-H) plot obtained from flow cytometry experiments using control and FN mutant MSCs. The control cell population is represented in blue, FN*_C123R_* mutant in green, and FN*_C231W_* mutant in red.

Previous work from us and others demonstrated impaired secretion of mutant FN in HEK293 cells and skin fibroblasts [16–18]. Therefore, we analyzed the secretion and assembly of FN in the iPSC-based model system by immunostaining. All stem cells were cultured in serum-free conditions; hence, the detected FN exclusively represented the cellular FN isoforms. Immunostaining for FN in control MSCs and chondrocyte pellets showed the expected FN matrix fibers (**Figure 2B**). However, a significant reduction in the assembled FN was noted in differentiated mutant MSCs and chondrocytes (**Figure 2B**). No alterations of FN mRNA levels were observed between control and mutant MSCs (**Figure 2C**). Immunoblotting of intracellular FN from cell lysates and secreted FN from culture media of control and mutant MSCs showed intracellular FN accumulation in the mutant cells with reduced levels of secreted FN outside the cells (**Figure 2D-F**). No evident FN degradation bands were noted in the cell lysate and culture medium. We also tested circulating plasma FN (pFN) levels in the patients by immunoblotting blood serum samples compared to an unaffected individual (**Figure 2G**). Both patient serum samples showed reduced pFN serum levels. Together, the data demonstrate that FN mutations leading to SMDCF decrease cellular and plasma FN secretion in differentiated MSCs and chondrocytes. Since FN promotes cell proliferation, we next analyzed the number of MSCs positive for the proliferative marker Ki-67 (**Figure 2H,I**). We observed for FN*_C123R_* and FN*_C231W_* a significant reduction of Ki-67 positive MSCs (**Figure 2H,I**). We also analyzed the forward scatter versus side scatter spectra of MSCs by flow cytometry (**Figure 2J**). The mutant MSC groups had higher side scatter distributions than the control MSCs. This means that the FN mutants have increased intracellular complexity, leading to more light scattering, presumably by intracellularly accumulated FN.

To obtain insight into the intracellular organization of the retained FN, we performed transmission electron microscopy (TEM) on control and mutant MSCs (**Figure 3A-C**). The control MSCs showed a typical organization and structure of cellular organelles such as mitochondria, rough endoplasmic reticulum (RER), and the Golgi apparatus (**Figure 3AI-III)**. In contrast, FN*_C123R_* and FN*_C231W_* exhibited numerous dense intracellular vesicles (**Figure 3B,C; arrowheads)**. These vesicles were coated with particles of ∼20-30 nm in size consistent with the size of ribosomes, suggesting that they originate from the RER and retain the mutant FN (**Figure 3B,C; arrowheads).** The extended RER frequently bulged out, apparently forming the vesicles that contained the dense material (**Figure 3BIII** **and CIII; arrowheads)**. Quantification of the vesicles showed a wide size range using the vesicle area on the TEM sections as a readout. They were between 0.04-30.9 µm^2^ for FN*_C123R_* with an average of 0.52 µm^2^ and between 0.08-70.7 µm^2^ for FN*_C231W_* with an average of 1.49 µm^2^ (**Figure 3D,E**). 59.2% of all vesicles in FN*_C123R_* MSCs and 31.3% in FN*_C231W_*MSCs were up to 2.5 µm^2^, whereas the remaining larger vesicles showed a broad size distribution. FN*_C231W_* MSCs were characterized by larger vesicles compared to FN*_C123R_*, and in some instances, the dense structures lost their vesicular-like conformation and expanded to over 50% of the entire cell (**Figure 3CI; black arrowheads)**. Presumably, the smaller vesicles fuse over time, giving rise to larger vesicles. The presence of ribosomes around these vesicles and the numerous RER-like projections extending into these vesicles suggest that the mutant FN is translocated into these dense vesicle-like structures originating from the RER.

**Figure 3.**
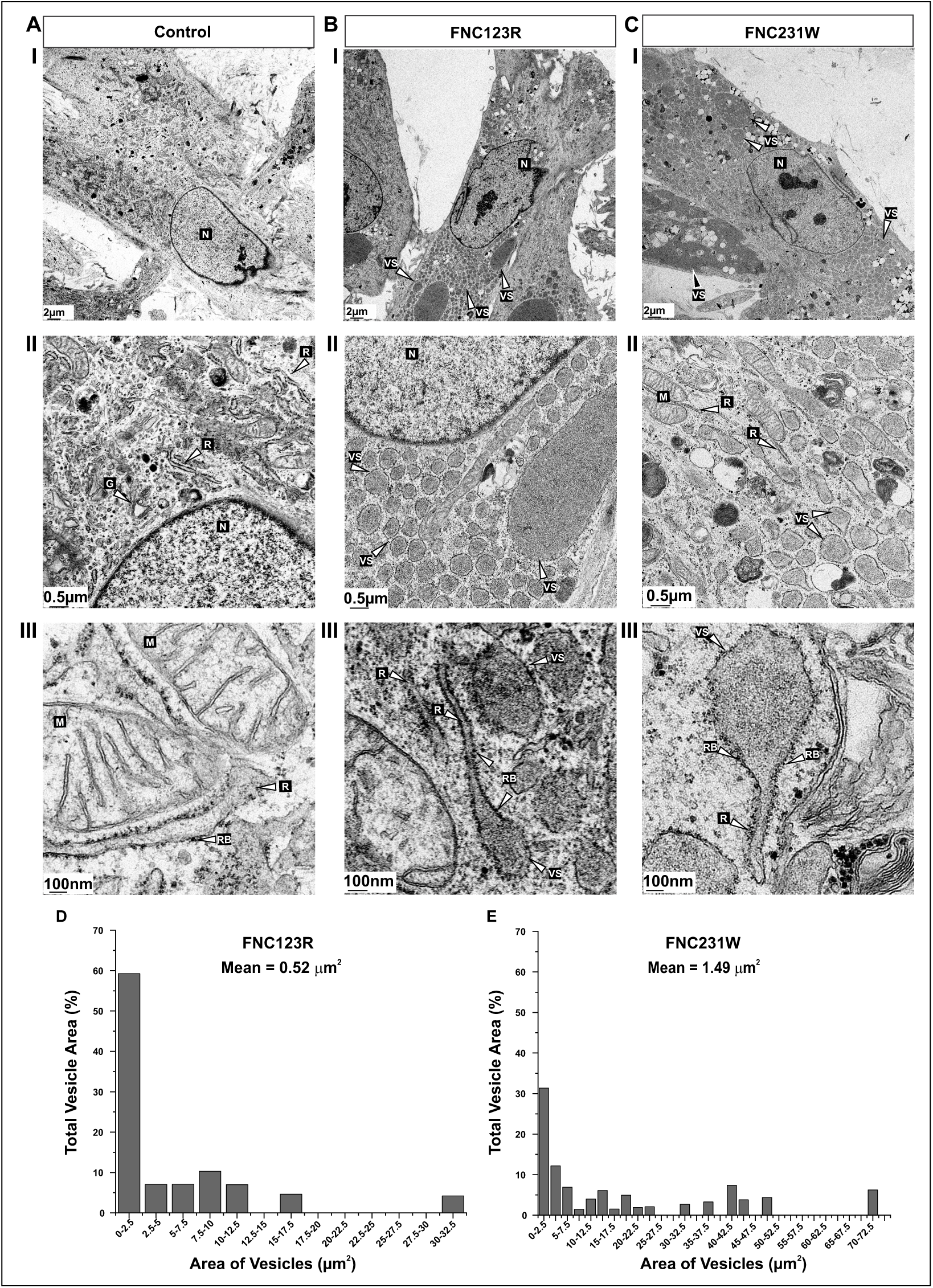
Transmission electron microscopy show massive accumulation of aberrant vesicular structures in the SMDCF MSCs. Transmission electron microscopy of **(A)** control, **(B)** FN*_C123R_*, and **(C)** FN_C231W_ MSCs. Magnifications are 1,200× in I, 6,800× in II, and 30,000× in III. **(A, I-III)** Control cells show normal nucleus (N), Golgi apparatus (G), rough endoplasmic reticulum (R), ribosomes (RB), and mitochondria (M) (arrowheads). **(B,C, I-III)** Patient MSCs show an accumulation of aberrant dense vesicular structures (VS) frequently covered with ribosomes (arrowheads). Arrowheads in III indicate vesicles bulging out from the rough ER in FN*_C123R_* and FN*_C231W_* MSCs. **(D-E)** Histograms of mutant protein-containing vesicle areas (bin size = 2.5 µm^2^). 100% represents the total area of all analyzed vesicles per mutant group. The mean vesicle area for each mutant is indicated. More than 700 vesicles were quantified per mutant group in a total of 10-12 cells.

To determine whether the vesicles in mutant cells contained FN, we performed 12 nm immunogold labeling of FN in control and FN mutant MSCs (**Figure 4**). In control cells, we observed FN labeling only in extracellular spaces on FN fibers but not within the cells (**Figure 4A; arrowheads)**. In the two FN mutant cells, the gold particles were prominently and uniformly present within the numerous intracellular vesicles of all sizes (**Figure 4B,C; arrowheads)**. The gold particles were irregularly distributed without any ordered pattern. The controls without the primary antibody showed the absence of gold labels, validating the specificity of the antibody against FN (**Figure 4A-C; left panel).** These data demonstrate that FN is intracellularly retained in the SMDCF mutant cells in numerous vesicular-like structures accumulating in the cytoplasm.

**Figure 4.**
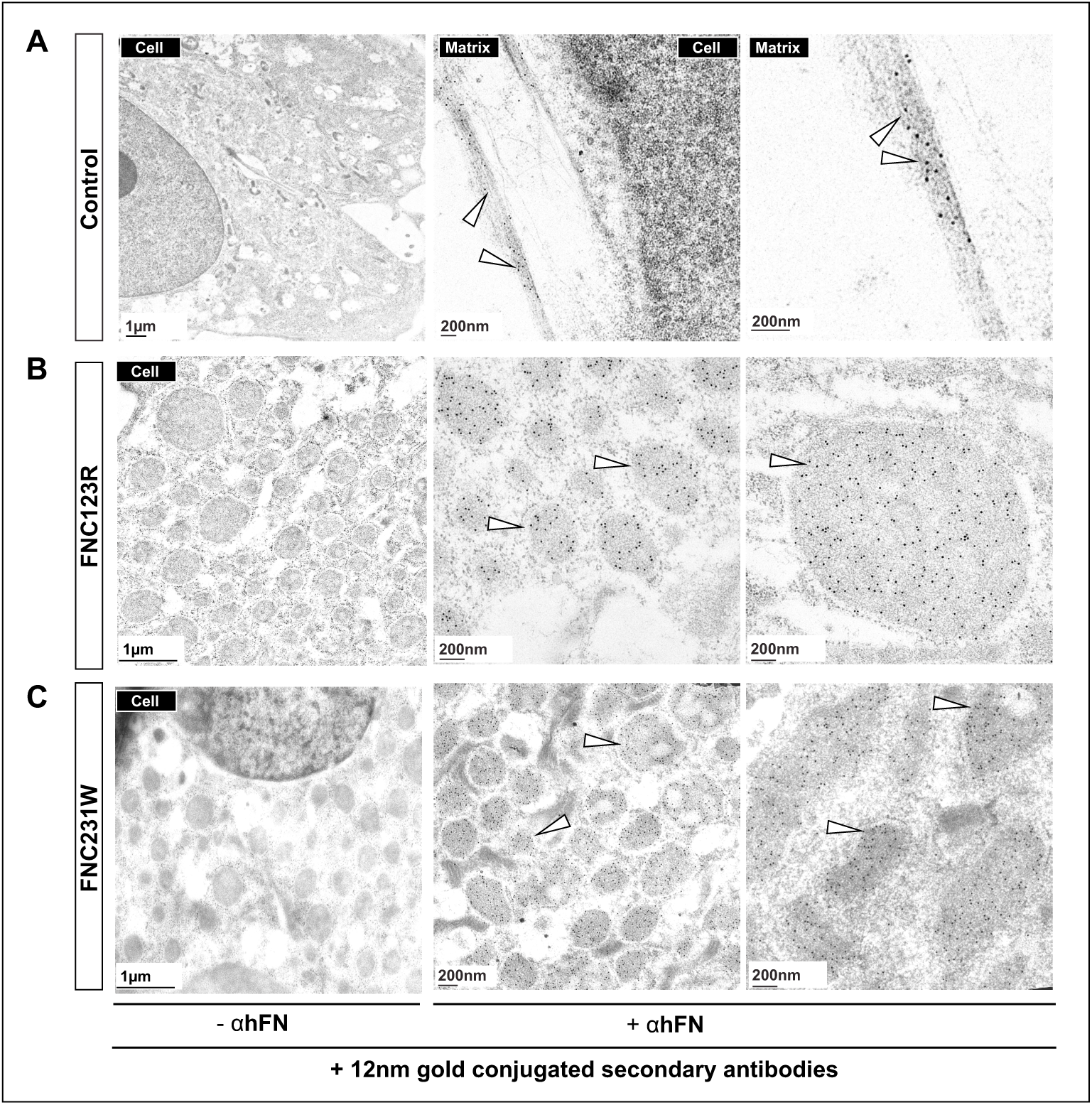
FN immunogold labeling of control and mutant MSCs. **(A-C)** Immunogold labeling of control and mutant MSCs was performed using an anti-FN antibody (+αhFN) and 12nm gold-conjugated secondary antibodies. **(A)** Control cells primarily showed gold particles on FN-positive fibers outside the cells (arrowheads). **(B-C)** FN mutant MSCs showed intensive gold labeling in all dense vesicular structures, confirming that the mutant FN accumulated in these aberrant vesicles (arrowheads). **(A-C)** The controls for each cell group without primary antibody (-αhFN) did not display any gold labels. Scale bars represent 1 µm.

### FN is exported directly from the rough endoplasmic reticulum to lysosomes in mutant cells

Since the TEM data showed that the vesicles in FN mutants are covered by ribosomes and appear to arise from the RER, we tested a series of ER and organelle markers in the MSCs using structured illumination microscopy [23,24]. We first analyzed the abundant ER chaperone BIP/GRP78, a member of the heat shock protein 70 family, which assists in protein folding and mediates the retention of misfolded proteins [25,26]. In control MSCs, we observed a punctate BIP staining pattern within the cells (**Figure 5A**). However, in the two FN mutant MSCs, a much stronger BIP staining was observed primarily in the FN-containing vesicles (**Figure 5A,B**). FN-positive vesicles in FN*_C123R_*and FN*_C231W_* MSCs were also positive for another ER-resident chaperone, HSP47, with significantly higher HSP47 levels per cell (**Figure 5C,D**). Interestingly, while all vesicles were positive for FN, small vesicles typically stained stronger than large vesicles, suggesting dynamic changes in FN distribution and/or organization as vesicles transition from small to large structures. The presence of ribosomes on the surface of the vesicular structures was validated by staining with antibodies against the 40S ribosomal subunit marker RPS6 [27]. Control MSCs displayed a dispersed ribosomal staining throughout the cytoplasm. The mutant cells, on the other hand, showed predominant localization of the RPS6 around the surface of the FN-containing vesicles (**Figure 5E**). The FN-containing vesicles in mutants were negative for early (RAB5) and late (RAB7) endocytic vesicle markers (**Figure 5F**) [28,29]. They were also negative for the autophagy marker LC3 and the Golgi marker GM103 (**Figure 5F**). However, the FN-accumulating vesicles were positive for lysosome-associated membrane protein 1 (LAMP1), and both mutants showed higher levels of LAMP1 per cell (**Figure 5G,H**). We confirmed these findings by analyzing another lysosomal marker, ATG5 [30], which demonstrated a similar staining pattern of the vesicles (**Figure 5I**). We also conducted high-resolution double immunogold labeling using 12 and 18 nm gold-conjugated secondary antibodies to examine the colocalization of the vesicle markers HSP47, BIP, and LAMP1 with FN in control and FN*_C123R_*MSCs (**Figure 6 A-C**). In the control cells, we observed scarce labeling of HSP47 and BIP near ER-like structures, whereas LAMP1 and FN were not detectable (**Figure 6A**). These findings aligned with the single FN labeling experiments shown in Figure 4. Conversely, in FN mutant MSCs, we detected dense gold labels marking FN in all vesicles together with gold particles labeling HSP47, BIP, and LAMP1 (**Figure 6B**). No nonspecific gold particles were present when the primary antibodies were omitted (**Figure 6C)**. These data are consistent with the light microscopic immunostaining data shown in Figures 5A,C,G. Altogether, these data suggest that in the FN mutant cells, misfolded FN, along with ER chaperones, are exported directly from the RER into variable-size vesicles budding out of the RER that are marked with ribosomes. These vesicles eventually transition into lysosomes, presumably to degrade the accumulating mutant FN.

**Figure 5.**
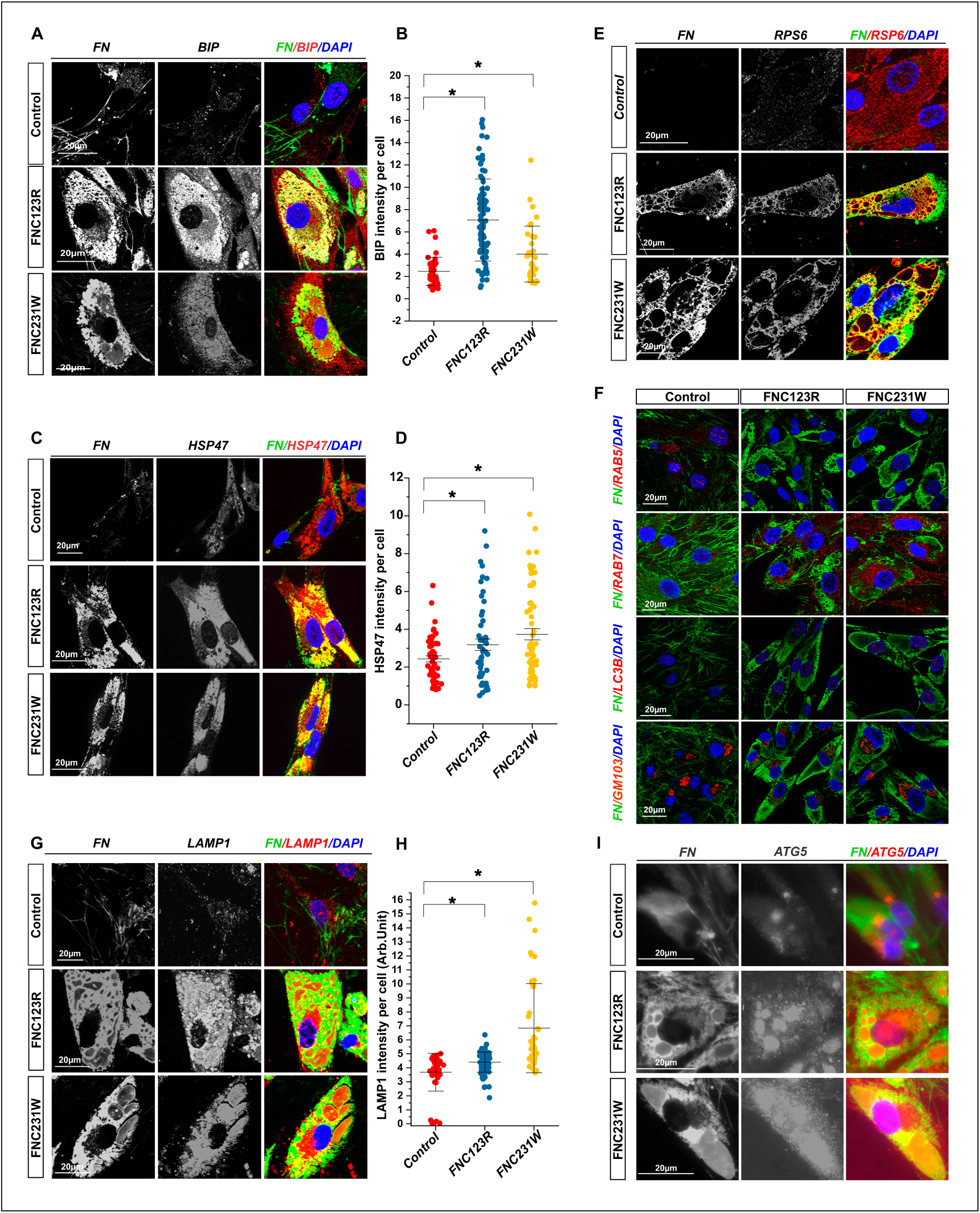
Indirect immunofluorescence analyses of subcellular compartment markers show that mutant FN is transported directly from the RER to lysosomes. Double immunostaining of control and mutant MSCs was performed with antibodies against FN (green) and against marker proteins for various subcellular compartments (red). The nuclear counterstain by DAPI appears in blue. Single channels are shown in grayscale, and overlaid images are pseudo-colored as indicated. Markers include the chaperones BIP **(A,B)** and HSP47 **(C,D),** the ribosomal marker RSP6 **(E)**, the early endosomal marker RAB5, late endosomal marker RAB7, autophagy marker LC3, and Golgi apparatus marker GM103 **(F)**, and the lysosomal markers LAMP1 **(G,H)** and ATG5 **(I)**. Figures display representative images of a total of three independent experiments (n=3). Each data point in the quantification graphs represents an individual cell, n = 41–121 cells per group. Error bars represent ± SD. Statistics were assessed by the unpaired Student’s two-tailed t-test with Welch’s correction and test for equal variance (F-test). Scale bars represent 20 μm for all images.

**Figure 6.**
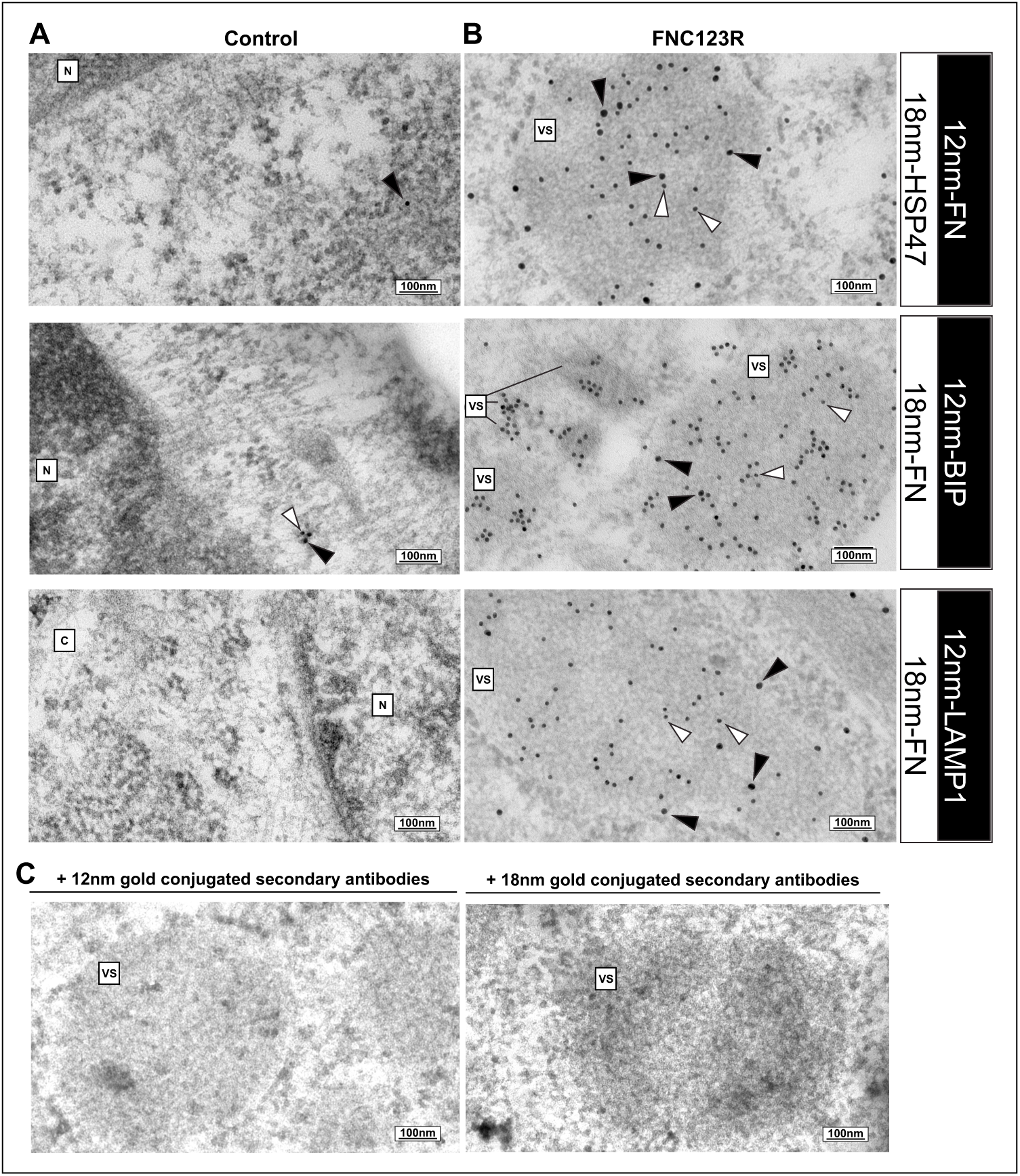
Double immunogold labeling of aberrant vesicles. **(A-B)** Double immunogold labeling of control and mutant MSCs was performed using anti-FN antibody (αhFN) with either ER markers BIP or HSP47 or with lysosomal marker LAMP1, using 12nm and 18nm gold-conjugated secondary antibodies. **(A)** Control cells showed very few gold particles for ER markers within the cells near the nucleus (arrowheads), but no label for LAMP1. **(B)** FN mutant MSCs showed double gold labeling in all dense vesicular structures for FN with the ER markers BIP or HSP47 and for FN with the lysosomal marker LAMP1 (arrowheads). **(C)** Controls without primary antibodies (only secondary antibodies) for each cell group did not display any gold labels. Scale bars are indicated in each image. Nucleus (N), Cytosol (C), Vesicular structures (VS).

### FN mutants exhibit transcriptomic dysregulation during chondrogenesis

To understand whether the increasing accumulation of the aberrant vesicles within the mutant cells and the lack of FN in the mutant cell matrix affected the overall transcriptome, we performed bulk RNA sequencing analysis (RNAseq) of the control and FN mutants at the MSC stage as well as after 21 d chondrogenic differentiation **(Supp.** Figure 2). Comparison of differentiated control chondrocytes to control MSCs showed more upregulated (2549) than downregulated (1359) genes **(Supp.** Figure 2A). This is expected as cells require a distinct cellular machinery to facilitate differentiation [31]. In contrast, comparing the FN mutant cells at the MSC stage or the differentiated chondrocyte stage with the respective control cells demonstrated a consistent pattern of more downregulated than upregulated genes **(Supp.** Figure 2B,C). At the MSC stage, 514-850 more genes were downregulated in the mutant versus control cells, which increased to 2112-2159 at the chondrocyte stage. This data suggests that the overall gene expression machinery progressively became downregulated during the experimental differentiation procedure of the two mutants, possibly due to the failure of the cells to undergo complete chondrogenic differentiation.

We then analyzed subsets of gene groups to identify the cellular processes altered in the mutants. We first analyzed HSP groups (*HSP40, HSP70, HSP90*) of protein folding chaperones, and we noted the most pronounced effects in the HSP40 gene family with a significant increase in 18-19 genes for FN*_C123R_* and FN*_C231W_* compared to control chondrocytes (**Figure 7A-C**). The HSP40 chaperone family is critical for facilitating protein folding, unfolding, and degradation [32], and the significant increase in many family members is likely a response to rescue the accumulation/misfolding of mutant FN in the SMDCF cells. We continued the analysis of the RNAseq dataset of the two mutant cell groups, focusing on some critical cellular stress markers (**Figure 7D**). At the MSC stage, FN*_C123R_* and FN*_C231W_* showed increased ER stress marker ATF6 and Calnexin levels. Several additional relevant genes were significantly upregulated upon differentiation into chondrocytes. These include HSPA5, the gene for BIP (see also Figure 5A), and EIF2A, a protein translation initiator factor associated with cellular stress response [33]. The mutant chondrocytes also exhibited increased expression of the ER degradation enhancing α-mannosidase-like proteins, *EDEM 2* and *3,* key players in facilitating ER-associated degradation of misfolded proteins [34,35]. Since the two analyzed FN mutations affect cysteine residues and disrupt a disulfide bond in FN type I domains, leading to a reactive free cysteine residue, we analyzed the protein disulfide isomerase gene family **(Supp.** Figure 3A-C). FN*_C123R_* and FN*_C231W_* chondrocytes showed significant dysregulation in several (5-12) of the disulfide isomerase enzymes, which was more pronounced for the FN*_C123R_* mutant (**Supp.** Figure 3B,C). Since we observed ribosomes on the aberrant vesicles, we analyzed the small and large ribosomal subunit markers (**Figure 8A-C**). Even though the global gene count was remarkably downregulated in the mutant cells, we observed a significant upregulation of the two ribosomal subunit genes (8-30 genes for the large subunit and 10-22 genes for the small subunit) in FN*_C123R_* and FN*_C231W_* chondrocytes (**Figure 8B,C**). This suggests that the mutant cells synthesize more ribosomal proteins to support the formation of numerous ribosomes that cover the FN-containing aberrant vesicles. These RNAseq data demonstrate that the mutant FN in the SMDCF stem cells resulted in cellular stress responses and the attempt of cells to rectify the accumulated or misfolded FN by protein folding chaperones. The data also suggests that aberrant disulfide bonds of the mutant FN play a role in the pathogenetic sequence. These mechanisms require enhanced ribosome activity to synthesize the respective proteins and enzymes.

**Figure 7.**
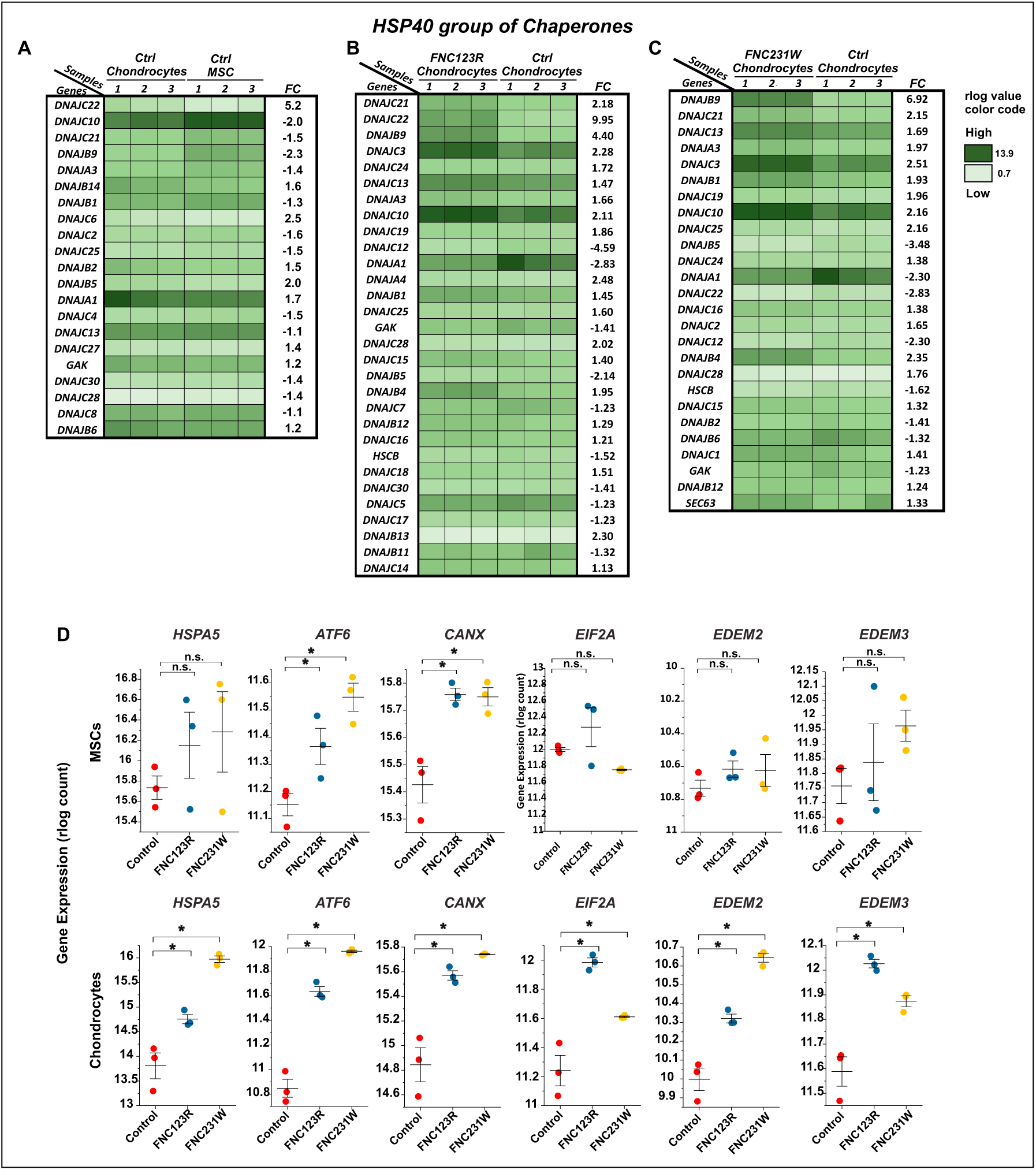
RNA sequencing analysis of protein folding chaperones and ribosomal markers. Control and mutant cells were analyzed by bulk RNA sequencing at the MSC and differentiated chondrocyte stage. **(A-C)** Heat maps of the differential gene expression for protein folding chaperones in the HSP40 group. The color codes for the rlog values are indicated on the right. FC represents fold-change. **(D)** Analysis of specific ER stress markers in control (red), FN*_C123R_* (blue), and FN*_C231W_* (yellow) MSCs (top panel) and chondrocytes (bottom panel). Statistical analysis was performed using nbinomWaldTest followed by the Benjamini-Hochberg method. For each group, genes with a significance of p ≤ 0.05 are shown in ascending order, with the lowest p-values on top.

**Figure 8.**
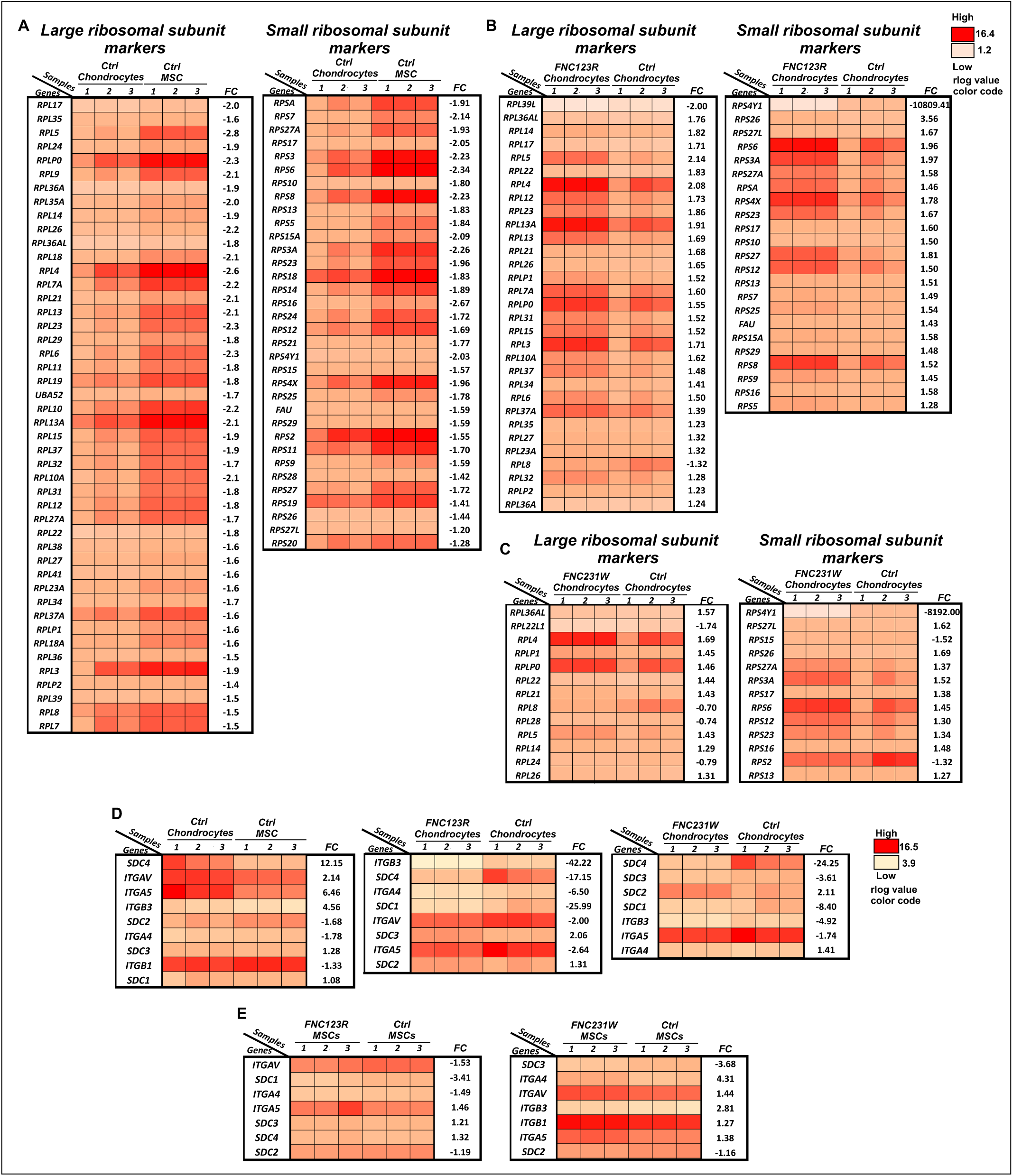
FN SMDCF mutant cells display increased ER stress markers and reduced expression of FN cell receptors. **(A-C)** Heat maps of gene expression are categorized into large and small ribosomal marker groups. For all heat maps, the rlog counts determined from 3 technical replicates per sample group are shown color coded. The color codes for the rlog values are indicated on the right. Statistical analysis was performed using nbinomWaldTest followed by the Benjamini-Hochberg method. For each group, genes with a significance of p ≤ 0.05 are shown in ascending order with the lowest p-values on top. **(D,E)** Heat maps of the mRNA levels of cell surface receptors for FN in chondrocytes **(D)** and in MSCs **(E)**. Gene expression is represented in rlog counts of 3 technical replicates per sample group. The color codes for the rlog counts are indicated on the right. The R package DESeq2 was used to identify differences in expression levels between the groups. Statistical analysis was performed using nbinomWaldTest followed by the Benjamini-Hochberg method. For the heat maps, genes with significance of p ≤ 0.05 are shown in ascending order with the lowest p-values on the top. FC represents the fold-change in rlog counts when comparing control chondrocytes to control MSCs **(D**, left panel), mutant chondrocytes to control chondrocytes **(D,** middle and right panels), and mutant MSCs to control MSCs **(E)**.

### FN mutant cells have reduced gene expression profiles for FN cell receptors

FN utilizes several integrins (e.g., α5β1, αvβ3, αvβ1, αvβ5, α4β1, α3β1) and heparan sulfate proteoglycans (e.g., syndecan (SDC) 2 and 4) as cell surface receptors that facilitate interaction with cells [36]. Assembly of FN depends on the interaction with integrins and syndecans, and these receptors have been previously reported to be essential for FN assembly [37–39]. Additionally, FN expression increases on the gene and protein level during chondrogenesis and assembled FN is essential for stem cell condensation [22]. Hence, we analyzed the expression level of relevant integrins and syndecans in the RNAseq dataset. Differentiated control chondrocytes versus undifferentiated MSCs showed a significant increase in SDC4 and all other pertinent SDCs and integrins, except ITGB1 and ITGA4 (**Figure 8D, left map)**. On the other hand, the two mutant FN*_C123R_* and FN*_C231W_* chondrocytes showed significant and consistent downregulation of most syndecans and integrins (**Figure 88D, middle and right maps)**. A similar downregulation of the two receptor groups was observed when mutant MSCs were compared to control MSCs (**Figure 8E**).

### FN mutants are characterized by altered mitochondrial structure

ER stress can lead to altered mitochondrial metabolism and function [40]. Since the FN mutants displayed elevated ER stress, we assessed the mitochondrial structure in control and mutant MSCs by TEM **(Supp.** Figure 4A). Control MSCs displayed typical mitochondria in size and distribution (**Supp.** Figure 4AI). In FN*_C123R_* and FN*_C231W_* MSCs, the mitochondria appeared damaged with less regular mitochondrial cisternae, and most of the mitochondria were elongated and swollen **(Supp.** Figure 4AII**,III)**. We continued analyzing the mitochondrial structure using the Mitotracker Deep Red FM dye **(Supp.** Figure 4B). Control MSCs displayed a typical perinuclear staining, whereas the mitochondria in the FN mutants appeared as elongated structures distributed throughout the cytoplasm (**Supp.** Figure 4B**; arrowheads).** We also noted an overall increase in cell size for FN*_C123R_* and FN*_C231W_* MSCs. The RNAseq dataset showed in control cells upon MSC to chondrocyte differentiation an overall downregulation of critical mitochondrial markers **(Supp.** Figure 3D). However, in the FN mutant cells, induction of chondrogenesis resulted in significant upregulation of mitochondrial markers relative to the control chondrocytes **(Supp.** Figure 3E,F). To evaluate if the structural changes in mitochondria of SMDCF cells also altered their function, we analyzed mitochondrial membrane potential and the ATP/ADP ratio **(Supp.** Figure 4C-E). However, these aspects did not change between mutant and control cells.

### Mutations in FN impair mesenchymal condensation and chondrogenesis

Mesenchymal stem cell condensation initiates chondrogenesis and induction of chondrogenic gene expression, promoting chondrocyte differentiation. Since FN is highly expressed during chondrogenesis, we investigated the consequences of FN mutations on this process in the iPSC-derived FN mutant stem cells using 3D micromass cultures (**Figure 9**) [41,42]. Condensation of control and FN mutant MSCs was determined microscopically on micromass 2D area projections (**Figure 9A,B**) [43–46]. Control MSCs began condensing within the first 12 h and formed a condensed micromass typically within 24 h. FN*_C123R_* and FN*_C231W_*MSCs, however, remained longer as large cell clusters and showed delayed micromass development (**Figure 9A**). Quantification at 6 d showed significantly larger areas for the mutant micromasses than for the control, confirming defects in MSC condensation (**Figure 9B**). To analyze differentiation into chondrocytes, we employed the conventional 3D pellet culture method for 21 d [42]. After chondrogenic differentiation, we analyzed the deposition of the chondrogenic marker collagen type II and FN by immunostaining (**Figure 9C,D**). Control micromasses showed high levels of collagen type II, which mostly co-localized with FN. On the contrary, the two mutant micromasses showed virtually no collagen type II deposition and a significantly reduced expression of the *COL2A1* mRNA (**Figure 9E**).

**Figure 9.**
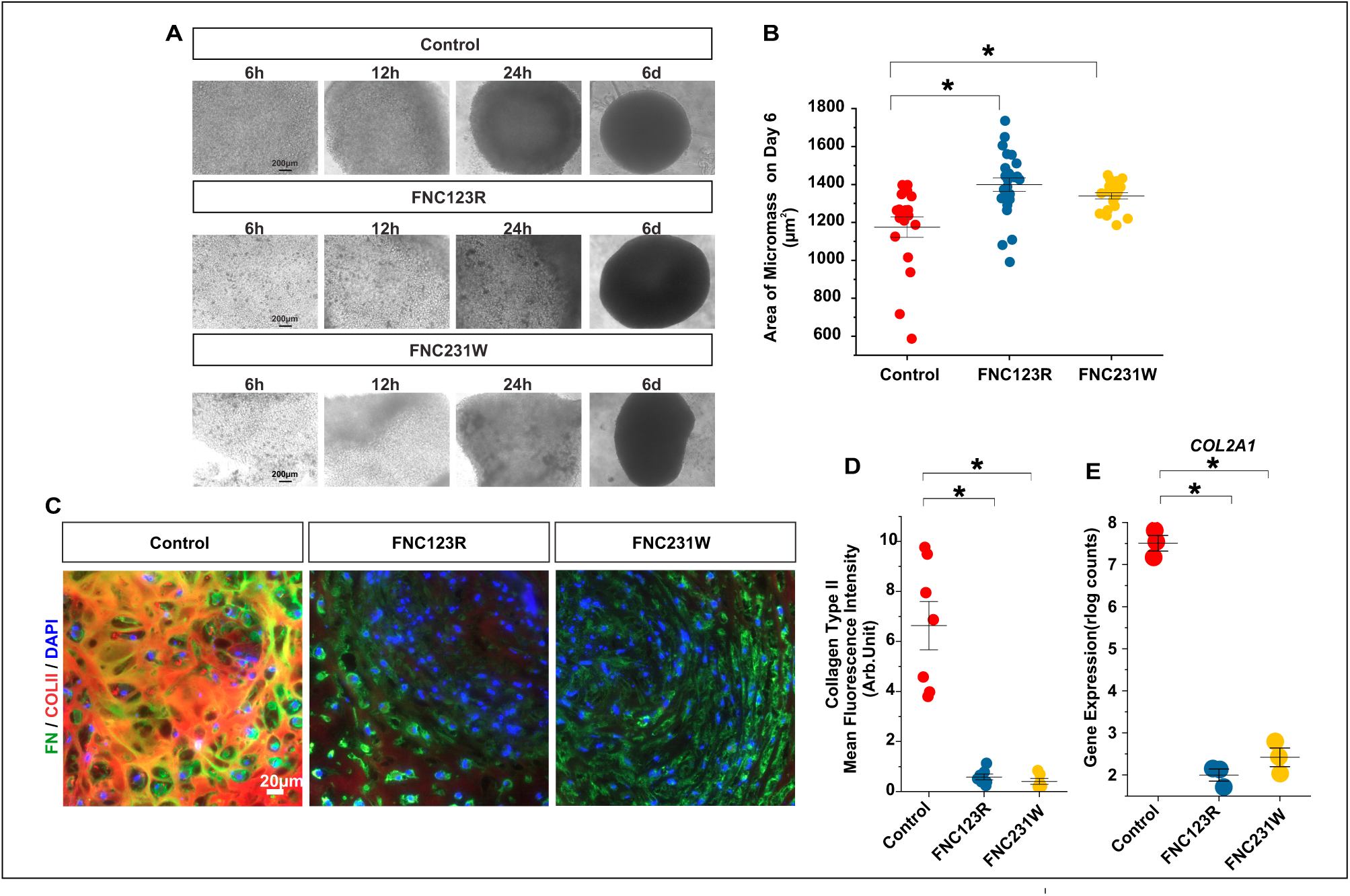
FN mutant cells exhibit impaired mesenchymal stem cell condensation and differentiation into chondrocytes. **(A)** Representative images of condensing MSCs in control and FN mutant cells from 6 h post cell seeding to 6 d. Scale bars represents 200 μm. **(B)** Quantification of micromass areas from images taken at 6 d for control, FN*_C123R,_* and FNC*_231W_* samples. The graph represents data from three independent experiments with a total n= 18-24 micromasses per sample group. Data are shown as means ± SEM. Statistical analysis was performed with the unpaired Student’s two-tailed t-test with Welch’s correction and test for equal variance (F-test). **(C)** Representative images of double immunostaining of FN (green) and collagen type II (red) in control and mutant differentiated micromass sections after 21 d chondrogenic differentiation. DAPI (blue) is used as a nuclear counterstain in all images. The scale bar represents 20 μm for all images. **(D)** Quantification of the mean fluorescence intensity of collagen type II in (C) from one representative experiment of a total of 3 independent experiments. The error bars represent ± SD. Statistical analysis was performed using one-way ANOVA with Bonferroni post-test. **(E)** *COL2A1* mRNA levels were obtained from the RNAseq data in differentiated control and mutant cells (n=3). Statistical analysis was performed using the nbinomWaldTest followed by the Benjamin-Hochberg method.

To further validate these findings, we analyzed the RNAseq data for other chondrogenic marker expression levels in the MSC and chondrocyte stages. We focused on proteoglycans, matrix markers, matrix metalloproteases (MMPs), cell surface markers, transcription factors, and growth factors **(Supp.** Figure 5A,B **and Figure 10A,B**). Differentiated control chondrocytes showed a consistent and significant increase in over 30 chondrogenic markers relative to undifferentiated control MSCs, confirming effective differentiation into chondrocytes **(Supp.** Figure 5A,B). On the contrary, most of those chondrogenic markers were significantly downregulated in FN*_C123R_* and FN*_C231W_* chondrocytes relative to the differentiated control chondrocytes (**Figure 10A,B)**. This data further substantiated the impaired chondrogenic differentiation capability of the two SMDCF mutant cell groups. Additionally, differentiated control chondrocytes showed increased expression of the FN gene (FN1), corroborating with the literature data **(Supp.** Figure 5A) [21,47]. However, mutant cells demonstrated a significant reduction in the FN expression level (**Figure 10A**). These data demonstrate that SMDCF mutations in FN impair the mesenchymal condensation and differentiation of MSCs into chondrocytes.

**Figure 10.**
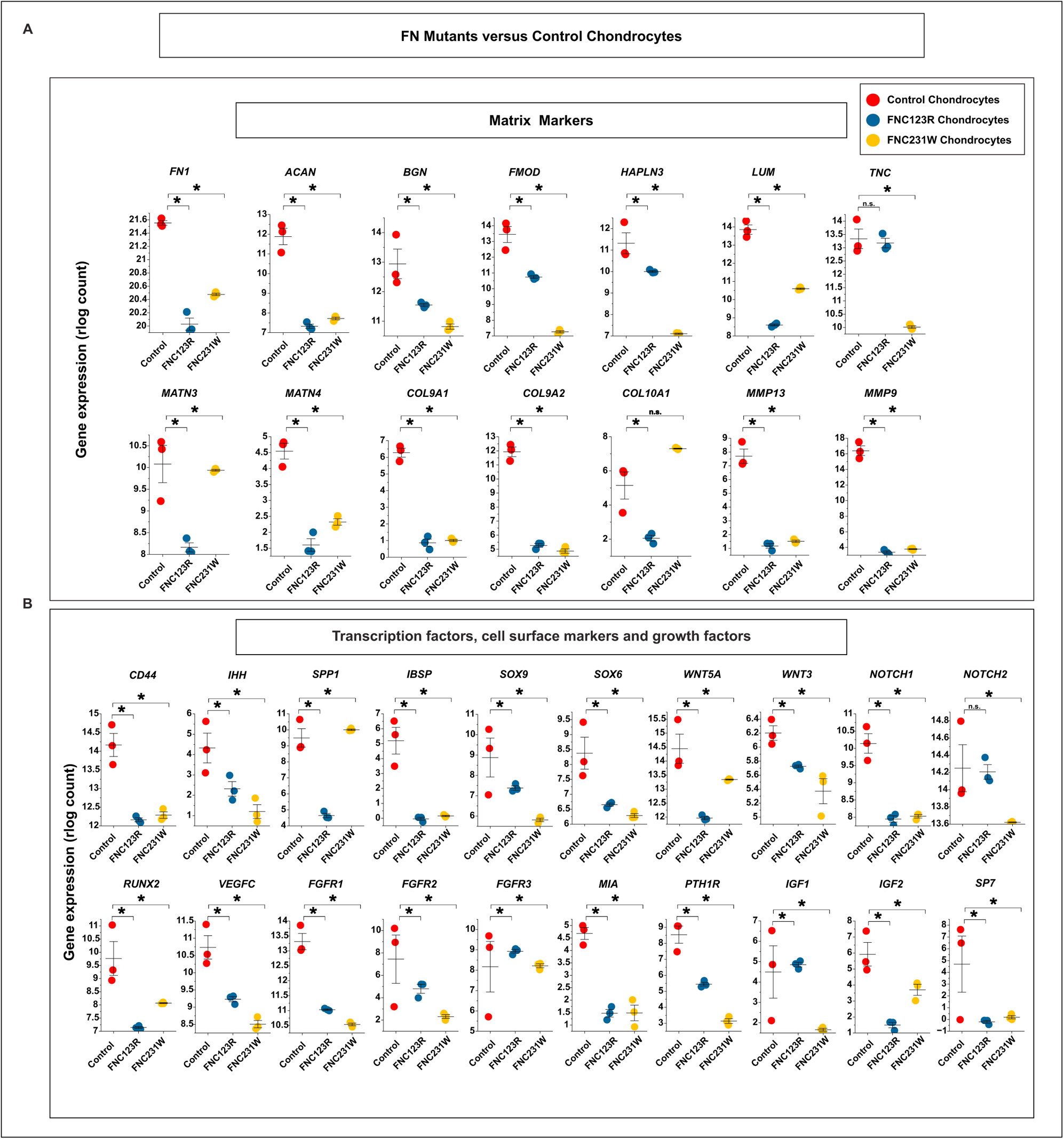
FN mutant cells exhibit reduced mRNA levels of chondrogenic markers. Control and mutant FN cells were analyzed by bulk RNAseq at the differentiated chondrocyte stage. Graphs represent selected key chondrogenic markers expressed in FN mutant cells versus control chondrocytes. The transcript reads for each marker are represented in rlog counts and sorted into two groups: **(A)** matrix markers and **(B)** transcription factors, cell surface markers, and growth factors. Data points represent 3 technical replicates per sample group. Statistical analysis was performed using the nbinomWaldTest followed by the Benjamini-Hochberg method. P ≤ 0.05 was taken as significant for all analyzed data.

### SMDCF mutations in FN1 alter the FN splicing pattern

The *FN1* gene gives rise to 27 transcripts by alternative splicing; 13 transcripts code for various FN isoforms, 1 is a non-coding transcript, and 13 are classified as retained introns **(Supp.** Figure 6A) [9]. Several cell culture studies explored the functional importance of some of these isoforms in chondrogenesis and skeletal development [7]; however, the exact physiological relevance and role in cartilage development remains unclear. Considering the importance of FN in stem cell differentiation and initiation of chondrogenesis, we analyzed the differential expression level of FN transcripts in the control and mutant cell RNAseq data. We examined the global transcript count of the 27 transcripts in the FN*_C123R_* and FN*_C231W_*mutant versus control cells in MSCs and differentiated chondrocytes and filtered those significantly dysregulated between the conditions **(Supp.** Figure 6B,C). Comparison of control chondrocytes and MSCs showed significant changes in 4 FN transcripts, including the coding transcript FN1-204 (V) and three retained intron transcripts **(Supp.** Figure 6B). Upon chondrogenic differentiation, the two FN mutants showed dysregulation of 7-9 protein-coding transcripts, most of which contained one or more of the alternatively spliced V, EDA, or EDB exons **(Supp.** Figure 6B). Significant changes in the same transcripts were noted for FN*_C123R_* MSCs versus the control MSCs **(Supp.** Figure 6C). However, the FN*_C231W_* MSCs exhibited alterations in only two *FN1* retained intron transcripts (FN1-224 and FN1-225) **(Supp.** Figure 6C). Analysis of differential expression of *FN1* transcripts in osteoarthritis (OA) lesion cartilage was recently published, demonstrating that OA is characterized by changes in the expression of specific *FN1* transcripts linked to reduced glycosaminoglycans and collagen type II expression [48]. We noted an overlap for some of these *FN1* transcripts between this study and our work (FN1-204,-209,-212,-224,-225, and-227). This suggests that the splicing and expression of specific *FN1* transcripts change during cartilage-associated pathologies.

### Exogenous treatment with FN and TGFβ1 can rescue impaired mesenchymal condensation of FN mutant stem cells

We next analyzed in the RNAseq dataset markers in chondrogenesis-regulating pathways that are potentially dysregulated in the FN mutants. We focused on three relevant groups: i) TGFβ and bone morphogenetic protein (BMP) signaling, ii) fibroblast growth factor signaling, and iii) Indian hedgehog signaling. We noted pronounced effects in the TGFβ/BMP group (**Figure 11A,B**). TGFβ1 was significantly elevated in control chondrocytes versus control MSCs and was dramatically downregulated in the mutant chondrocytes versus control chondrocytes (**Figure 11A,B**). TGFβ1 promotes mesenchymal condensation [49], and FN and TGFβ1 modulate each other in chondrogenesis [7]. Therefore, we focused on investigating the role of TGFβ1 in the SMDCF cells.

**Figure 11.**
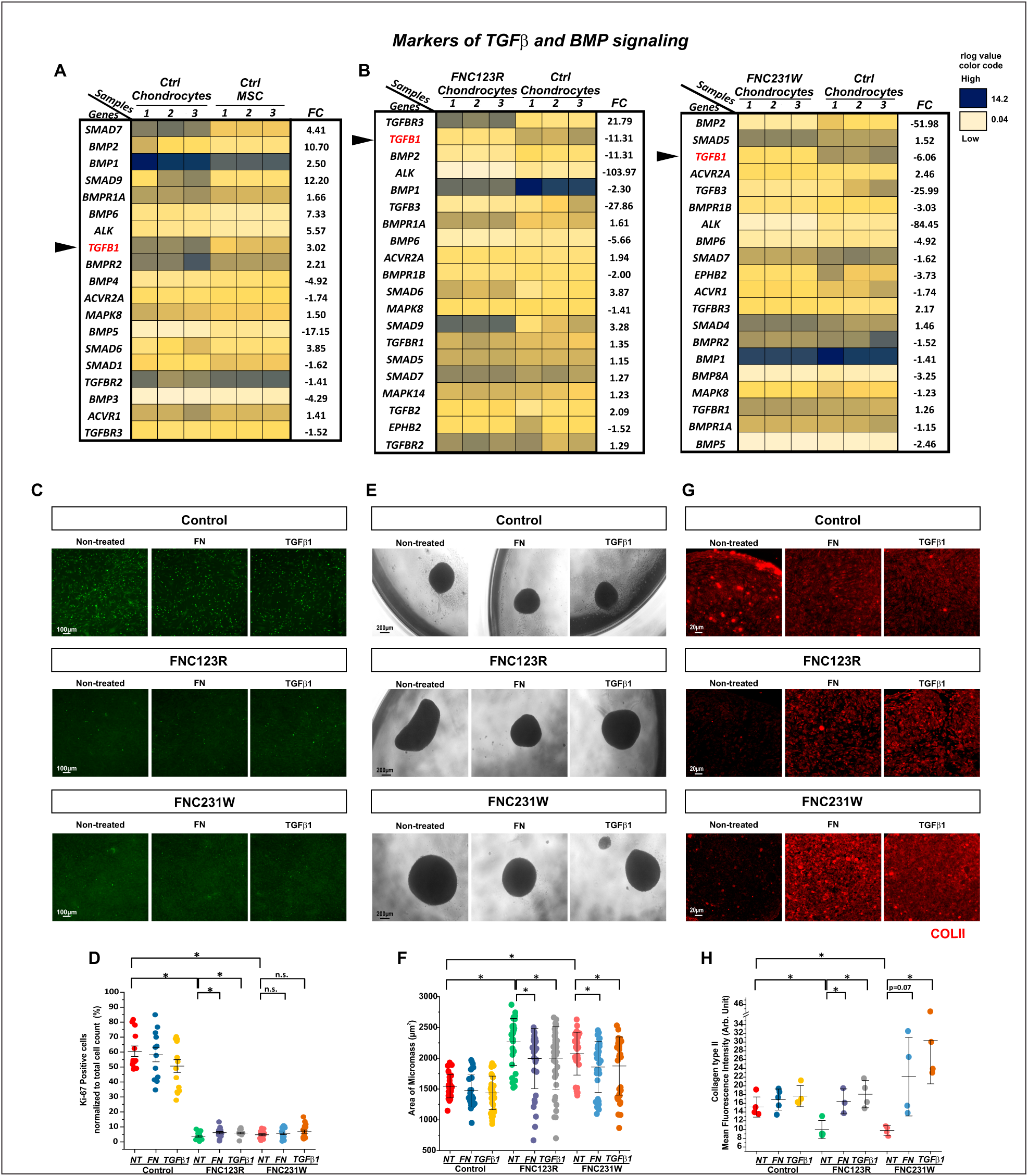
Exogenous supplementation of FN and TGFβ1 improves condensation defects in FN mutant cells. **(A,B)** Heat maps of RNAseq data for markers in the TGFβ and BMP signaling pathways comparing **(A)** control MSCs and control chondrocytes, and **(B)** control and FN mutant chondrocytes. The rlog counts were determined from 3 technical replicates per sample and group. Statistical analysis was performed using the nbinomWaldTest followed by the Benjamini-Hochberg method. For each group, only the mRNAs with a significance of p ≤ 0.05 are shown in ascending order, with the lowest p-values on the top. FC represents fold-change. **(C)** MSCs were analyzed for the presence of proliferation marker Ki-67 (green) using immunostaining under non-treated and FN (25 μg/mL) or TGFβ1 (10 ng/mL) treated conditions. **(D)** Quantification of Ki-67 positive cells shown in (C) normalized to the total cell count. The graph represents Ki-67 positive cells from 12-17 images obtained from three independent experiments (n=3). Each data point represents the average percentage of Ki-67 positive cells from one individual image. The error bars represent ± SEM. Significance was evaluated by the unpaired Student’s two-tailed t-test with Welch’s correction and test for equal variance (F-test). **(E)** Representative images of the live cell condensation assay of control versus FN mutant cells in non-treated and in FN-or TGFβ1-treated conditions at 6 d. **(F)** Quantification of micromass areas shown in (E). The graph represents data from five independent experiments with a total n=30-40 micromasses per sample group. Data are shown as mean ± SEM. Statistical analysis was performed with the unpaired Student’s two-tailed t-test with Welch’s correction and test for equal variance (F-test). **(G)** Representative images of collagen type II immunostaining of control versus FN mutant cell micromass sections at 6 d either non-treated or treated with FN or TGFβ1. **(H)** Quantification of collagen type II immunostaining shown in (G). The graph represents the mean intensity of collagen type II from micromasses. Each data point represents the average mean fluorescence intensity from individual images per micromass with a total of n=3-5 micromasses per sample group. Data are shown as mean ± SD. Statistical analysis was performed with the unpaired Student’s two-tailed t-test.

Cell proliferation of stem cells and chondrocytes is essential for chondrogenic differentiation and long bone growth. Since we noted that the FN mutant MSCs stained significantly lower for the cell proliferation marker Ki67 (**Figure 11C,D**). Hence, we then investigated whether exogenous supplementation of purified FN (25 µg/mL) or recombinant human TGFβ1 (10 ng/mL) in the culture medium could improve the proliferation defect. FN*_C123R_* MSC showed a small increase in Ki67-positive cells upon treatment with either FN or TGFβ1 but not FN*_C231W_*MSCs (**Figure 11C,D**). Further, we analyzed if supplementation of FN or TGFβ1 could rescue the mesenchymal condensation defect in the FN mutants using 3D micromass cultures. No changes were observed in control stem cell condensation upon treatment with FN or TGFβ1. However, the FN*_C123R_*and FN*_C231W_* MSCs displayed a clearly visible and quantifiable improvement in mesenchymal condensation upon adding either FN or TGFβ1 (**Figure 11E,F**). Further, we analyzed these micromasses by collagen type II immunostaining. The level of collagen type II remained similar in control micromasses upon treatment with FN or TGFβ1 (**Figure 11G,H**). However, when FN*_C123R_* and FN*_C231W_* mutant cells were treated with exogenous FN or TGFβ1, collagen type II deposition was fully rescued to the control levels (**Figure 11G,H**).

As iPSC controls were limited at the beginning of the study, most of the experiments were performed with one biological control iPSC line. However, as iPSCs became commercially available, we validated this rescue experiment with additional sex-matched iPSC controls (Ctrl2 and Ctrl3). Ctrl2 and Ctrl3 iPSCs were cultured and validated for pluripotent and MSC markers upon differentiation (**Supp.** Figure 7A-D). Relative to these control cell groups, the FN*_C123R_* and FN*_C231W_* mutants displayed a significantly delayed MSC condensation, clearly visible by larger micromass sizes at 6 d **(Supp.** Figure 7E). Consistent with the initial results (**Figure 11E,F**), the condensation defect was markedly alleviated in the two mutants after exogenous supplementation of FN or TGFβ1, notable by a significant reduction in micromass sizes **(Supp.** Figure 7E). No alterations in condensation were observed in the control cell groups when treated with FN or TGFβ1. These findings demonstrate that the addition of FN or TGFβ1 can ameliorate the stem cell condensation defect in the SMDCF mutant cells.

## Discussion

FN is a vital regulator of various organ systems in embryonic and postnatal development. However, virtually nothing is known about the consequences of SMDCF-causing FN mutations on cell physiology and pathology. In the present study, we have developed an SMDCF model system using iPSCs generated from fibroblasts obtained from two previously reported SMDCF patients harboring FN mutations p.C123R and p.C231W in FNI domains at the N-terminal region of FN (16, 17). Since skeletal growth and development defects are the predominant clinical phenotype reported in individuals with SMDCF, we differentiated the SMDCF iPSCs into MSCs and subsequently into chondrocytes to investigate the consequence of FN mutations on stem cell differentiation and chondrogenesis.

In line with previous work [16], we show with the patient-derived SMDCF MSCs that the mutant FN protein is not secreted and accumulates within cells. Since these are heterozygous mutations, the expected relative amounts of FN dimers are 25% for wild-type and 25% for mutant homodimers, and 50% for wild-type/mutant heterodimers. However, our results show very little FN secreted by the mutant cells, suggesting that the wild-type FN containing homo-and heterodimers were co-retained together with mutant FN homo-and heterodimers in the patient cells. This initiated a cellular stress response, mitochondria structure alterations, and increased synthesis of protein folding chaperones such as BIP and HSP47. On the contrary, autophagy was not activated. Both analyzed mutations inevitably disrupt known disulfide bonds in the FNI-2 domain (pattern Cys1-Cys3) and the FNI-5 domain (pattern Cys2-Cys4) [16,17,50]. Almost all other known SMDCF-causing mutations also disrupt disulfide bonds in FNI domains [16–18]. AlphaFold predicted that both mutations introduce relatively minor structural changes within the mutation-containing domain as well as in directly adjacent FNI domains. This aligns with previous structure predictions of p.C87F [16]. The data suggests that consequences arising from a disrupted disulfide bond in an FNI domain are more relevant for the disease mechanism than any other gross structural changes within the individual FNI domains. One possible and likely consequence is that the uncoupled cysteine residue provides a reactive free sulfhydryl group that can form an aberrant disulfide bond either with other mutant or unaffected FN molecules or with other proteins harboring free cysteine sulfhydryl groups. Both scenarios are predicted to lead to protein aggregation and misfolding [51]. In addition, previous studies have reported that mutations of selected conserved residues (other than cysteines) in FNI-1 to 5 also impair FN secretion because these N-terminal FNI domains act as a unit to facilitate proper folding and secretion [3,52].

We noted in SMDCF MSCs numerous vesicles in the cytosol ranging from ∼100 nm – 2 µm in diameter. These vesicles contained FN and the chaperones BIP and HSP47 and had the signature of ribosomes covering the surface. Electron microscopy showed that these vesicles bud out directly from the RER. We interpret these results as a means for the cells to clear the RER after the chaperones failed to mediate proper FN folding. We consistently observed that the immunofluorescence light microscopy staining for FN was more intense in the smaller vesicles and relatively weaker within larger vesicles in mutant cells. This difference is likely attributed to variations in the distribution or aggregation of FN within these vesicles as they transition into larger structures. Many of the cytosolic vesicles were positive for the lysosomal markers LAMP1 and ATG5, suggesting a transition or fusion into lysosomal structures, presumably in an attempt to degrade the accumulating FN. These observations are consistent with ER-to-lysosome-associated degradation (ERLAD). Similar cellular mechanisms have been previously reported for several pathological conditions, including the α1-antitrypsin deficiency diseases [53,54]. Misfolded α1-antitrypsin that accumulates within the RER of liver cells is directly exported from the RER to lysosomes for ER-to-lysosome associated degradation [54,55]. Another example of ERLAD comes from experiments with hyper-stimulation of the thyroid, where thyrotrophic hormone-producing cells develop intracisternal granules in the RER, causing the RER to dilate and transition into cisternae containing the β subunits of thyrotrophic hormone [56]. These cisternae converted directly into lysosomes to facilitate protein degradation. Typically, BIP and other ER chaperones that escape the RER are retrieved in COPI vesicles via the KDEL receptor recognition system from post-ER compartments back to the ER lumen [57]. However, we observed the presence of chaperones together with FN in the vesicular structures in mutant SMDCF cells. Mutant FN aggregation or misfolding apparently triggered its export from the RER with associated chaperones bypassing the KDEL recognition system in the Golgi apparatus.

As the identified vesicles become lysosome-like structures, the expectation is that lysosomal proteases would degrade the accumulating FN. However, our immunoblotting analysis of total protein cell lysates from mutant cells revealed no visible FN degradation. This observation is further substantiated by the presence of numerous vesicles containing accumulated FN that persists within cells. This suggests that either the mutant protein complexes undergo very slow and minimal degradation or that aggregated or misfolded FN is densely packed and potentially cross-linked through additional disulfide bonds. This dense packing may hinder lysosomal proteases from docking onto their recognition sites, thereby preventing the degradation of mutant FN. This notion is consistent with our electron microscopy analyses, which demonstrate rich and dense labeling of the vesicles with anti-FN antibodies. Alternatively, it is possible that chaperones complexed with FN aggregates prevent lysosomal FN degradation. It will be necessary to further analyze the degradation mechanisms of mutant FN in these aberrant vesicles and to test strategies that potentially facilitate lysosomal FN degradation, for example, using lysosome-enhancing agents [58]. Interestingly, there are no reported defects in other organ systems of the patients with SMDCF, while the same pathological molecular pathways should be in place in other cells. For example, one would expect a liver phenotype, supported by our findings of reduced levels of circulating FN in FN*_C123R_* and FN*_C231W_* mutant cells, suggesting impaired FN secretion from hepatocytes. Since hepatocytes are specialized liver cells with an enhanced ability to clear toxins and misfolded proteins, it is possible that the degradation of misfolded mutant FN in those cells is more efficient, thereby preventing functional defects in the liver [59].

Cell proliferation and differentiation are closely associated during cartilage development. Hence, we analyzed the proliferation of iPSC-derived FN mutant MSCs, demonstrating a significant reduction of stem cell proliferation in the two mutant cell groups. This contrasts with a previous study, reporting the absence of proliferative effects using skin fibroblasts from an SMDCF patient with a p.C97W mutation in FN [18]. A possible explanation is that a FN deficient matrix and/or the abnormal protein accumulation in the SMDCF cells may impact stem cells differently than fibroblasts. The reduced proliferation potential of SMDCF MSCs likely plays an important role in the context of pathologic chondrogenesis, as mesenchymal progenitor cells normally have to undergo extensive proliferation to increase cell density and allow enhanced cell-cell contacts to facilitate mesenchymal condensation [60–62], and differentiation into chondrocytes [63]. FN promotes cellular proliferation through integrins in various cells, including chondrocytes [64]. Thus, the absence of FN in the ECM of SMDCF cells is likely a key contributing factor to the reduced proliferation observed in these cells. We attempted to rescue this defect with exogenous supplementation of purified FN to the mutant cells. This led to only a small improvement in cell proliferation for *FN_C123R_* but not for *FN_C231W_* cells. We suspect that this could be due to the reduced expression of key FN interacting integrins subunits (e.g., ITGB1, ITGB3, ITGAV) observed in the FN mutants, which i) are predicted to reduce the interaction of FN with cells and ii) may affect the assembly of exogenously provided FN.

We demonstrated that the FN mutant stem cells display delayed condensation and impaired chondrocyte differentiation marked by a reduction in collagen type II at the mRNA and protein levels. This is in contrast to the recently identified osteoarthritis-causing FN mutation p.C518F in the FNI-8 domain, which did not impair FN secretion but reduced FN binding to collagen type II [65]. The fact that in SMDCF, the impaired FN secretion severely affects MSCs’ differentiation into chondrocytes and expression of collagen type II, as well as a broad spectrum of other chondrogenic markers, demonstrates the pathologic severity of the SMDCF mutations on cellular differentiation.

One of the possible reasons for the impaired chondrogenesis in SMDCF could be a reduced or altered interaction of FN with integrins or syndecans, as both regulate chondrogenesis and the expression of chondrogenic markers *in vivo* [14,66,67]. A deficiency of FN in the matrix and reduced expression of these surface receptors noted in the mutant cells could presumably disrupt FN-receptor interaction efficiency in SMDCF, dysregulating chondrogenesis. Alternatively, the impaired chondrogenesis could result from defects in TGFβ signaling, which is indispensable for cartilage development [49,68,69]. Both FN and TGFβ have intertwined roles in chondrogenesis [7]. FN increased TGFβ levels and signaling during chondrogenesis in several cell culture studies [70]. In turn, TGFβ also promoted the expression of FN and differential isoform expression in chondrocyte cultures [7]. In the present study, we identified that TGFβ1 mRNA was significantly downregulated in the FN mutant cells when subjected to chondrogenic differentiation. Whether the absence of FN in the matrix directly diminished TGFβ1 levels, first impaired stem cell differentiation resulting in reduced TGFβ1 expression, or whether the altered TGFβ1 level alone was the primary driver for defective chondrogenesis in the mutant cells remains to be established. Nevertheless, purified FN or active TGFβ1 supplementation partially rescued defective stem cell condensation and differentiation of the *FN_C123R_*and *FN_C231W_* mutant cells. Based on these cell culture data, supplementation with exogenous FN and TGFβ may be beneficial in the future to alleviate pathological aspects in SMDCF. However, testing this hypothesis is currently hampered by the lack of a suitable SMDCF mouse model.

In summary, the present study explores the molecular pathomechanisms underlying FN mutations that cause SMDCF, leading to the model outlined in **Figure 12**. In unaffected cells under normal physiological conditions (**Figure 12A**), FN is co-translationally translocated into the ER lumen, undergoing dimerization, post-translational modifications, and protein folding. Folded FN is sorted into transport vesicles that carry the protein from the RER to the Golgi apparatus and into secretory vesicles, which are targeted to the plasma membrane for release into the ECM. The secreted compact FN dimer interacts with cell surface receptors, triggering assembly into a fibrous FN network. The FN matrix promotes MSCs to undergo mesenchymal condensation and chondrogenic differentiation associated with increased levels of FN and TGFβ1, facilitating skeletal development. In cells harboring disease-causing SMDCF mutations (**Figure 12B**), FN and ER chaperones are intracellularly retained inside aberrant vesicles that originate from the rough ER and transition into lysosomes. These defects induce elevated cellular stress, altered mitochondrial structure, reduced cell proliferation, matrix deficiency in FN, and reduced levels of TGFβ1 upon chondrogenic induction. These molecular defects affect the mutant MSCs to differentiate into chondrocytes, contributing to skeletal dysplasia development characteristic for SMDCF.

**Figure 12.**
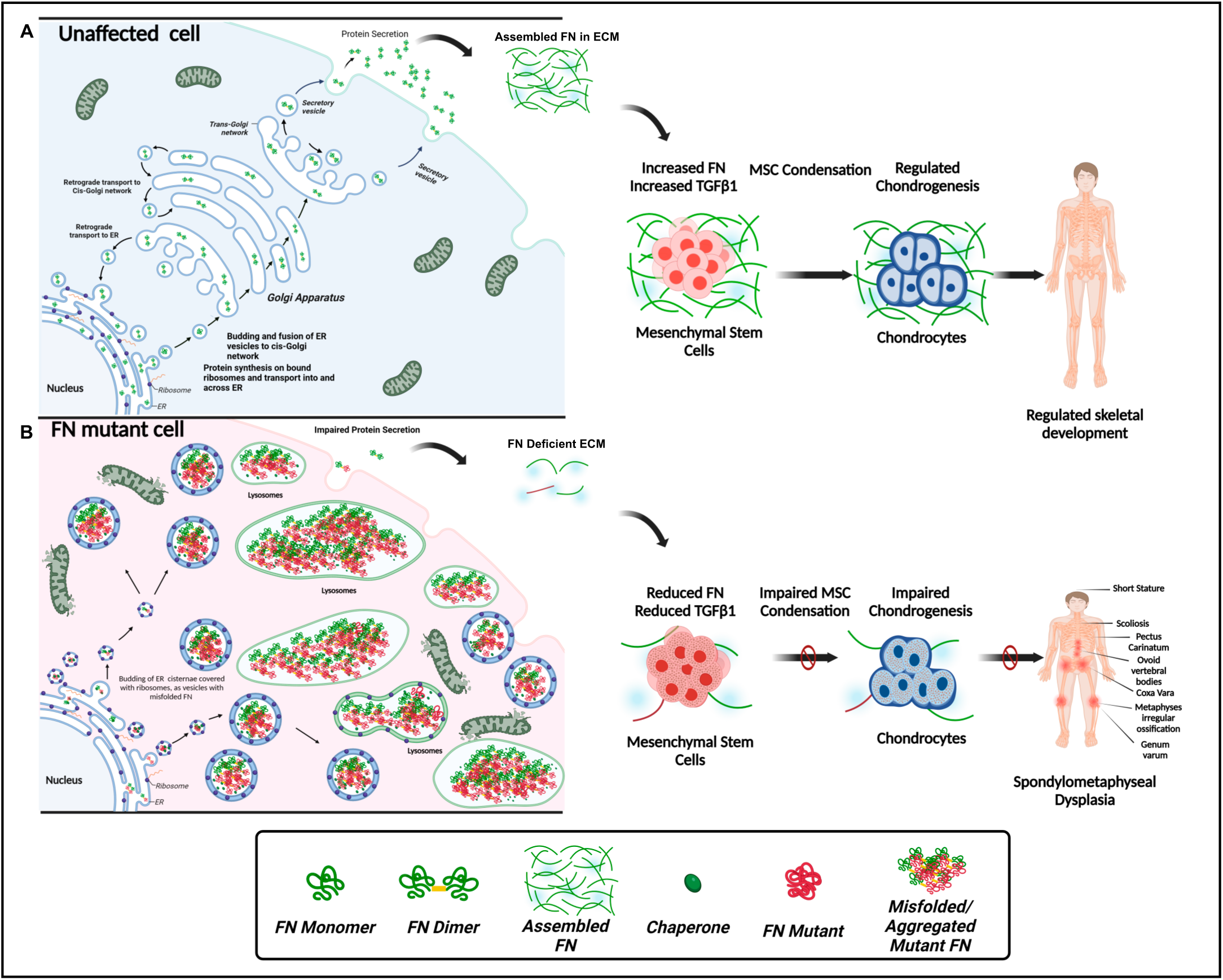
Schematic representation of the molecular mechanism and pathways proposed for SMDCF pathogenesis. **(A)** Schematic overview of FN secretion and assembly by an unaffected cell and extracellular FN fibers in mesenchymal condensation and chondrogenesis. **(B)** Overview of the proposed intracellular and extracellular mechanisms contributing to the pathogenesis of SMDCF in FN mutant stem cells. Details are described in the main text. The various components in the figure are not drawn to scale.

## Materials and Methods

### IPSC Culture and Differentiation into Mesenchymal Stem Cells

Skin fibroblasts from SMDCF patients with FN mutations FN*_C123R_* and FN*_C231W_* and from an unaffected individual were used to generate induced pluripotent stem cells (IPSCs) at the core facility of CHU-Saint Justine (Montreal, Canada). Fibroblasts were reprogrammed to iPSCs with integration-free Sendai virus expressing Oct4, Sox2, Klf4, and c-Myc, using CytoTuneTM-iPSC reprograming kit (Life Technologies; Catalogue #A13780-01, A13780-02), followed by validation of the iPSC colonies by staining with antibodies for markers SSEA-4, Sox2, OCT4, and TRA1, as described previously [71]. Generation and use of the patient cells was approved by Institutional Review Board (IRB) (Protocol number: <u>MP-21-216-962, 4181</u>). Additional control iPSCs were purchased from the Coriell Institute for Medical Research (GM23338 and GM23716). IPSCs were cultured using a feeder free method with E8 serum free essential medium (Life Technologies; Catalogue #A1517001), supplemented with antibiotic Primocin (100 µg/mL, Invivogen; Catalogue #ant-pm-2), and 2mM L-Glutamine (Stem Cell Technologies; Catalogue #07100;). Geltrex Matrix (800 µg/mL, Gibco; Catalogue #A14133-0), a reduced growth factor basement membrane matrix, was used as the coating substrate for iPSC cultures. Cells were treated with E8 medium supplemented with rock inhibitor RevitaCell supplement (Gibco; Catalogue #A2644501) for 18-24 h for post thaw recovery of cryopreserved iPSCs and during passaging. Gentle Cell Dissociation Reagent (Stem Cell Technologies; Catalogue #07174) was used for passaging the colonies. IPSCs were validated for the expression of pluripotent markers using flow cytometry outlined below.

IPSCs were differentiated into MSCs using the STEMdiff Mesenchymal Progenitor Kit (Stem Cell Technologies; Catalogue #05240;). The differentiation protocol involved treatment of IPSCs with STEMdiff Animal Component-Free (ACF) Mesenchymal Induction Medium for 4 d, followed by expansion and further culturing with MesenCult ACF medium (Stem Cell Technologies; Catalogue #05445), supplemented with 2mM L-Glutamine (Stem Cell Technologies; Catalogue #07100) for 21 d. The mesenchymal progenitor cells were tested and validated for the expression of MSC markers (see below). All MSCs were cultured on ACF Cell Attachment Substrate coated surfaces (Stem Cell Technologies; Catalogue #07130). ACF Cell Dissociation Kit (Stem Cell Technologies; Catalogue #05426) was used for passaging the MSCs for expansion and experiments. All stem cells were cultured at 37°C, 5% CO_2_, 5%O_2,_ and 90% humidity.

### Differentiation and Analysis of Mesenchymal Stem Cells in Chondrocytes

MSCs were differentiated into chondrocytes using the MesenCult-ACF Chondrogenic Differentiation Kit (Stem Cell Technologies; Catalogue #05426) comprising of MesenCult-ACF Chondrogenic Differentiation Basal Medium (Stem Cell Technologies; Catalogue #05456) and MesenCult-ACF 20× Chondrogenic Differentiation Supplement (Stem Cell Technologies; Catalogue #05457). The chondrogenic induction medium was supplemented with the 1× differentiation supplement and 2mM L-Glutamine. MSCs were differentiated either by 3D micromass culturing for 6 d for the cell condensation assay (1×10^6^ cells/mL) or alternatively by 3D pellet culturing for 22 d (5×10^5^ cells per micromass) in 15mL conical tubes (Corning; Catalogue #352096). All cells were cultured at 37°C, 5% CO_2_, 5%O_2,_ and 90% humidity. Following differentiation, micromasses or pellets were either fixed with 4% paraformaldehyde (PFA) for 24 h or used directly for RNA and protein extraction. Fixed masses were paraffin-embedded and sectioned at 4μm thickness for immunostaining and histological analysis.

### Cell Condensation Assay

MSCs were cultured and differentiated into chondrocytes by 3D spheroid/micromass culturing technique using the Nunclon Sphera 96 well plates (Thermo Fisher Scientific; Catalogue #174927) that prevented cell adhesion to the plate surface and promoted spheroid formation for live cell condensation assays. For micromasses, cells were seeded at high density and as small microliter droplets onto culture plates. 20µL of a 1×10^6^/mL cell suspension were plated per well and incubated for 1 h, followed by addition of 150µL chondrogenic induction medium. Condensation of micromasses was imaged from 1 h to 6 d using a light microscope (Axiovert 40CFL) and the ZEN 2 (blue edition) software. Rescue experiments were performed by supplementing the medium with either 25 µg/mL human plasma fibronectin (Millipore Sigma; Catalogue #FC010) or with 10 ng/mL of human TGFβ1 (Peprotech; Catalogue#100-21). At the experimental end point at 6 d, post imaging the samples were fixed with 4% PFA for 1 h followed by paraffin embedding and sectioning (4μm) for immunostaining and histological analysis. Images of micromasses were used for quantification of total micromass area using Image J software [72]

### Flow Cytometry

IPSCs were immunostained for pluripotent markers C-MYC, NANOG, OCT4 and SOX2 and MSCs were stained for CD73, CD105 and an undifferentiated IPSC marker TRA-1-60 using specific antibodies conjugated with fluorophores (Supp. Table 1). Cell density in each sample was maintained at 5×10^5^ cells/mL for all analysis. Antibody labeling and flow cytometric analysis followed standard procedures using the BD LSR Fortessa 5L flow cytometer and the BD FACS Diva 8.0.2 software at the Advanced BioImaging Facility of the McGill Life Sciences Complex (Montreal, Canada). The FlowJo 10.8.1 software was used for further data analysis.

### Transmission Electron Microscopy (TEM) and Immunogold Labeling

For TEM, MSCs were seeded at a density of 75,000 cells/well on Nunc Lab-Tek 8-well chamber permanox slides (Thermo Fischer Scientific; Catalogue #177445) and cultured for 3 d. Cells were fixed with 2.5% glutaraldehyde in 0.1 M cacodylate buffer at 4°C overnight, washed with 0.1 M cacodylate washing buffer, post-fixed with 1% aqueous osmium tetroxide and 1.5% potassium ferrocyanide for 1 h at 4°C, and en bloc stained with 2% uranyl acetate for 1 h. After fixation, the cells were dehydrated with increasing ethyl alcohol, embedded with Epon, and polymerized at 60°C for 48 h. The blocks were sectioned using a Leica EM UC7 ultramicrotome (Leica Microsystems, Wetzlar, Germany) with a Diatome diamond knife (Diatome, Nidau, Switzerland), and the ultrathin sections (70-80nm) transferred onto 200-mesh Cu TEM grids. The sections were counterstained with 4% uranyl acetate and Reynold’s lead. For immunogold labeling experiments, cells were fixed with 4% PFA including 0.05% glutaraldehyde in 0.1M sodium cacodylate buffer for 1 h and then embedded in LR white resin, followed by UV light-mediated polymerization in a Leica EM AFS2 embedding system. Sections were placed on formvar coated mesh copper grids and used for either single or double immunogold labeling with anti-human fibronectin (polyclonal and monoclonal), BIP (polyclonal), HSP47 (monoclonal) and LAMP1 (polyclonal) primary antibodies. 12 nm colloidal gold AffiniPure Donkey anti-rabbit antibody and 18 nm colloidal gold AffiniPure Goat anti-mouse antibody were used as secondary antibodies (Supp. Table 1). Counterstaining was performed as described before with 4% uranyl acetate and Reynold’s lead. Imaging was performed on a FEI Tecnai 12 120kV TEM microscope at the McGill University Facility for Electron Microscopy Research Facility using a AMTV601 CCD camera.

### Sanger Sequencing and qPCR

Cells were seeded at a density of 1.5×10^5^ cells/well in 6 well plates (Corning; Catalogue #3516) and cultured for 3 d. Total RNA was extracted using the RNeasy kit (Qiagen; Catalogue #74104) as per the manufacturer’s instructions. cDNA was prepared with 0.1-1 µg of total RNA using the Protoscript First Strand cDNA Synthesis kit (New England Biolabs; Catalogue #E6560L) as per the manufacturer’s instructions. For real-time qPCR, experiments were performed using an SYBR Select Master Mix (Applied Biosystems, Catalogue# LS4472908). For sanger sequencing PCR reactions were performed using specific oligos against the target FN domains (Supp. Table 2) and Taq DNA Polymerase kit (ZmTech Scientifique; Catalogue #T207025) and analyzed on 2% agarose gels. The amplified PCR products were extracted from the gel using the QIAquick Gel Extraction Kit as per manufacturer’s instruction (Qiagen; Catalogue #28704), followed by Sanger DNA sequencing of the PCR products at the Génome Québec (Montreal, Canada).

### RNA Sequencing and Data Analysis

MSCs were cultured for 10 d (1.5×10^5^ cells in triplicates per cell group) and chondrocyte differentiation was performed for 21 d as described above. RNA was extracted using the RNeasy kit (Qiagen; Catalogue #74104) as per the manufacturer’s instructions. RNA samples (150 ng/µL) were analyzed at the McGill Genome Centre (Montreal, Canada) for quality and integrity (RIN) using the Qubit RNA High Sensitivity Assay and TapeStation. All samples were found to have high RIN values between 8.2-9.5. Ribosomal RNA was depleted, and cDNA libraries were prepared using 1 µg RNA per sample. Bulk RNA sequencing was performed with the Illumina NovaSeq6000 system. Processing of the raw RNAseq dataset and differential gene expression analysis was performed by the Canadian Center for Computational Genomics (C3G; Montreal, Canada). The Ensemble GRCh38.p13 (Genome Reference Consortium Human Build 38) was used as the reference database for alignment of the reads. GenPipes is the main in-house framework of the C3G used to perform major processing steps [73]. Adaptor sequences and low-quality score-containing bases (Phred score < 30) were trimmed from reads using Trimmomatic [74]. The resulting reads were aligned to the genome, using STAR [75]. Read counts were obtained using HTSeq [76]. The R package DESeq2 was used to identify differences in expression levels between the groups using negative Binomial GLM fitting and Wald statistics (nbinomWaldTest) [77]. The R package *ashr* was used to shrink log2 fold changes in gene expression data [78]. All data sets in figures represent gene expression in regularized log counts (rlog counts) unless specified otherwise in the figure legends. For gene group sets, the HUGO Gene Nomenclature Committee (HGNC) classification was used as a reference.

### In silico Structure Prediction

Bioinformatic structure predictions were performed using the ColabFold v1.5.3: AlphaFold2 using MMseqs2, a simplified version of AlphaFold v2.3.1 [79]. Sequence for human fibronectin was obtained from the Uniprot database (Sequence ID: P02751-15, correlating to DNA reference NM_212482.4). Structure predictions were made for the FN domain type I domains 1, 2, 3, 4 and 5 using the respective sequences from P02751-15. To predict structural consequences of the FN mutations FN*_C123R_* and FN*_C231W_*were introduced into the wild-type FN sequences. For both wild-type and FN mutant domains, structures with highest ranking were analyzed and selected. Structure visualization and alignment was performed using the PyMOL software [80,81]. Root mean square deviation (RMSD) for alignment of mutant domain structures with the respective wild-type domains were obtained using the CE algorithm in PyMOL.

### Immunoblotting

For immunoblotting analysis, cells were lysed using RIPA buffer (50 mM Tris, pH 8.0, 150 mM NaCl, 0.5% sodium deoxycholate, 1% Triton X-100, 0.1% sodium dodecyl sulfate) supplemented with 2% v/v protease inhibitor cocktail (Roche; Catalogue #11697498001) and 1% v/v phosphatase inhibitor cocktail (Sigma-Aldrich, Catalogue #P5726). Lysates were centrifuged at 180 × g for 5 min to pellet the debris and the clear lysates were used for immunoblotting. Total protein concentration was determined by the BCA protein assay kit (Thermo Fisher Scientific; Catalogue #23225). Total protein ranging from 20-30µg was used in a standard immunoblotting procedure. All antibodies used in various experiments with the respective dilutions are listed in Supp. Table 1. Horseradish peroxidase-conjugated goat anti-rabbit, or anti-mouse secondary antibodies were incubated for 2 h at ambient temperature. Super signal chemiluminescent western blotting substrate (Thermo Fisher Scientific; Catalogue #34580) was used for the development of blots, and images were recorded using the ChemiDoc MP imaging system (Bio-Rad). Band intensities were quantified with ImageJ [72] and normalized to the GAPDH control.

### Analysis of Human Plasma Samples

Human plasma samples were obtained from the two SMDCF patients with the FN mutations p.C123R and p.C231W, and from an unaffected individual (parent of patient with the p.C123R mutation). Plasma samples were analyzed for fibronectin by immunoblotting using 0.1 µL plasma sample under reducing conditions with a specific anti-human FN antibody (Supp. Table 1).

### Immunostaining and Immunofluorescence Microscopy with Apotome

For indirect immunofluorescence staining, cells were plated on Nunc Lab-Tek 8-well chamber permanox slides (Thermo Fischer Scientific; Catalogue #177445) at a density of 75,000 cells per well and cultured as specified in the figure legends. Cells were fixed with 4% PFA or 4% PFA including 0.1% Triton X-100, depending on the experimental setup, blocked with 10% v/v goat serum, and incubated overnight at 4 °C with either one or two primary antibodies (raised in different species) for single or double immunostaining, respectively (Supp. Table 1). Alexa Fluor 488 goat anti-rabbit antibody, Cy3 goat anti-mouse antibody, or Cy5 goat anti-mouse antibody were used in combination or alone as secondary antibodies (Supp. Table 1). Nuclei were counter-stained with 4, 6-diamidino-2-phenylindole (DAPI) (Abcam; Catalogue #188804). An epifluorescence microscope (Zeiss; AxioImager M2) with an Orca Flash 4.0 CMOS camera was used for analysis. All images were originally in grayscale and then pseudo-colored using the Zen Pro software version 2.6 (Zeiss). Superresolution Structured Illumination Microscopy was performed using the Zeiss Apotome 2 fitted with the AxioImager M2. Apotome calibration and validation of the imaging technique were performed using 0.2μm TetraSpeck microspheres (Invitrogen; Catalogue #T7280) and Zeiss immunofluorescence calibration slides (Zeiss; Catalogue #000000-1213-943). Post processing of apotome images included phase correction and deconvolution.

### Image Quantifications

Quantification of the mean fluorescence intensity of specific markers was performed by processing of the images as described previously in detail [82]. Mean intensity per cell were quantified by outlining single cell area. Quantification of Ki-67 positive cells were performed using a procedure described previously [82]. The number of Ki-67 positive cells was normalized to the total number of DAPI stained cells per image. Quantification of the area of micro masses in cell condensation assays were performed by outlining the micromasses using the Wacom Intuos Graphic Tablet with Stylus (Model no. CTH490AK) in ImageJ. Similarly, for quantification of area and number of intracellular vesicles from the EM micrographs, vesicles were outlined using the freehand tool in Image J with the Wacom tablet and stylus, and the data was plotted as a histogram using the “distribution” statistical function in the Origin Pro software.

### Statistics

All data are represented as means ± standard deviation (SD) or standard error of the mean (SEM), depending on the experimental setup as indicated in the figure legends. Experiments were performed either as triplicates or quadruplets. Significance was evaluated using either Student’s two-tailed t-test, or one-way or two-way ANOVA post Bonferroni test as specified in each figure legend. All statistical analyses were performed using the OriginPro version 2023 software (OriginLab). Outlier analysis was performed using the Grubb’s test with confidence levels set to 95%. Data with p-values ≤0.05 were considered significant and are denoted as *. Non-significance is labeled “n.s.”.

### Schematic Figures

All schematics in this manuscript were prepared using the Biorender software (URL: https://app.biorender.com/)

## Data Availability

The RNA sequencing data generated in the study has been submitted to the NCBI’s Gene Expression Omnibus database (https://www.ncbi.nlm.nih.gov/geo) and is available through GEO accession number GSE251698. Key resources used in this study are provided in Supplemental Tables 1 and 2.

## Acknowledgments

We express our sincere gratitude to the SMDCF patients for donating samples for this research study. Additionally, we would like to acknowledge the Facility for Electron Microscopy Research at McGill University, for their assistance with access and preparation of samples for electron microscopy. We are also grateful to the McGill Genome Centre for performing the RNA sequencing and to the Canadian Center for Computational Genomics for their assistance with the RNAseq data analysis.

## Author Contributions

NEHD, PMC, and DPR conceived the study, designed the experiments, analyzed the data, and wrote the manuscript (with critical input from all authors). NEHD performed the experiments. JR, MK, and JM provided various methodological contributions. DFM provided reagents and critical inputs for the study. NEHD, PMC, and DPR acquired the funding.

