## Supplemental Data and Information for "Mutations in Fibronectin Dysregulate Chondrogenesis in Skeletal Dysplasia"

#### Supplemental Methods

***Mitotracker and Mitochondrial Membrane Potential Analysis***

Cells were seeded at a density of 75,000 cells per well in 8-well chamber slides for immunofluorescence imaging and in 12-well plates for flow cytometry analysis and cultured for 3 d. For immunofluorescence analysis, the cells were incubated after washing with buffer containing 200nM Mitotracker Deep Red Dye (Invitrogen; Catalogue #M22426) for 15 min at 37$^{o}$C, fixed with 4% PFA for 3 min, and counterstained with DAPI. Immunofluorescence imaging was performed on fixed samples using a Zeiss AxioImager M2 and Zen Software version 2.6. For flow cytometry analysis, cells were detached from wells using ACF gentle dissociation reagent and centrifuged at 180 × g for 5 min. The pellet was resuspended in PBS containing 200 nM Mitotracker dye CM-$H_2X_{ROS}$ (Invitrogen; Catalogue #M7513) and incubated for 15 min at 37$^{o}$C. The cells were then fixed with 4% PFA and analyzed by flow cytometry at the Advanced BioImaging Facility of the McGill Life Sciences Complex, McGill University (Montreal, Canada) using the BD LSR Fortessa 5L flow cytometer and the BD FACS Diva 8.0.2 software.

***ATP/ADP Ratio Assay***

ATP/ADP ratio assay was performed with the differentiated MSC cells using the luminometry ATP/ADP Ratio Assay Kit (Sigma-Aldrich; Catalogue #MAK135-1KT). Cells were seeded at a density of 10,000 cells/well in solid white 96-well microplates (Greiner Bio; Catalogue #655073) and grown for 5 d, washed, and analyzed per manufacturer's instructions. Luminescence readings were recorded using a PerkinElmer 2030 (Model # 2030-0020).

### Supplemental Figure 1

**A**

SMDCF and Control iPSC Information

| iPSC Name | Domain Containing Mutation | Age | Sex | Ethnicity |
|---|---|---|---|---|
| FNC123R | FNI-2 | 4 years | Female | African origin |
| FNC231W | FNI-5 | 13 years and 9 months | Male | Vietnamese origin |
| Control | Not Applicable | 41 years | Male | NA |
| GM23338 | Not Applicable | 55 years | Male | White American origin |
| GM23716 | Not Applicable | Fetal | Female | African origin |

**B**

| Mutation | Domain containing mutation | RMSD of affected domain | Neighboring domain | RMSD of neighboring domain |
|---|---|---|---|---|
| C87F | FNI-1 | 0.25Å | FNI-2 | 0.24Å |
| C169Y | FNI-3 | 0.61Å | FNI-2 | 0.05Å |
| C213Y | FNI-4 | 0.64Å | FNI-5 | 0.07Å |
| C225W | FNI-4 | 0.26Å | FNI-5 | 0.08Å |
| Y240D | FNI-5 | 0.26Å | FNI-4 | 0.15Å |
| C258Y | FNI-5 | 0.93Å | FNI-4 | 0.11Å |
| C260G | FNI-5 | 0.23Å | FNI-4 | 0.14Å |

**C**

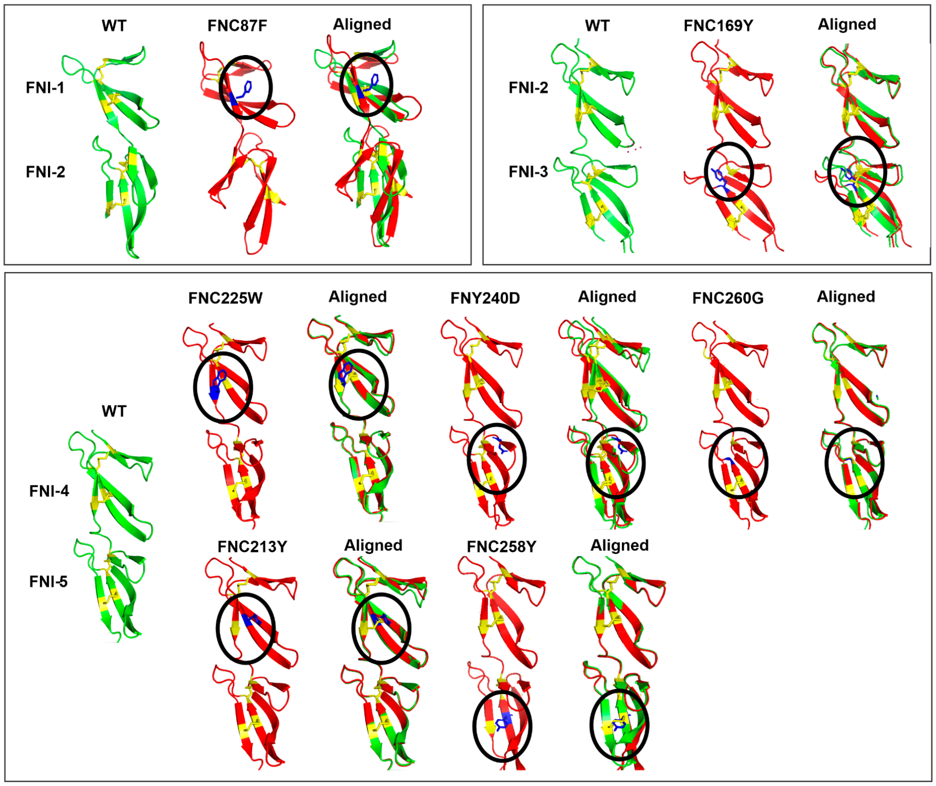


**Supp. Figure 1.** ***In silico*** **structure prediction of the FN mutant domains.**

**(A)** Overview of the donor details for the iPSC lines used in the study. **(B)** Summary of the RMSD values from structure alignments of previously reported SMDCF-causing mutations in FN. **(C)** Overview of the predicted FN mutant structures, aligned with the corresponding wild-type FN domains using AlphaFold2 and PyMOL.

#### Supplemental Figure 2

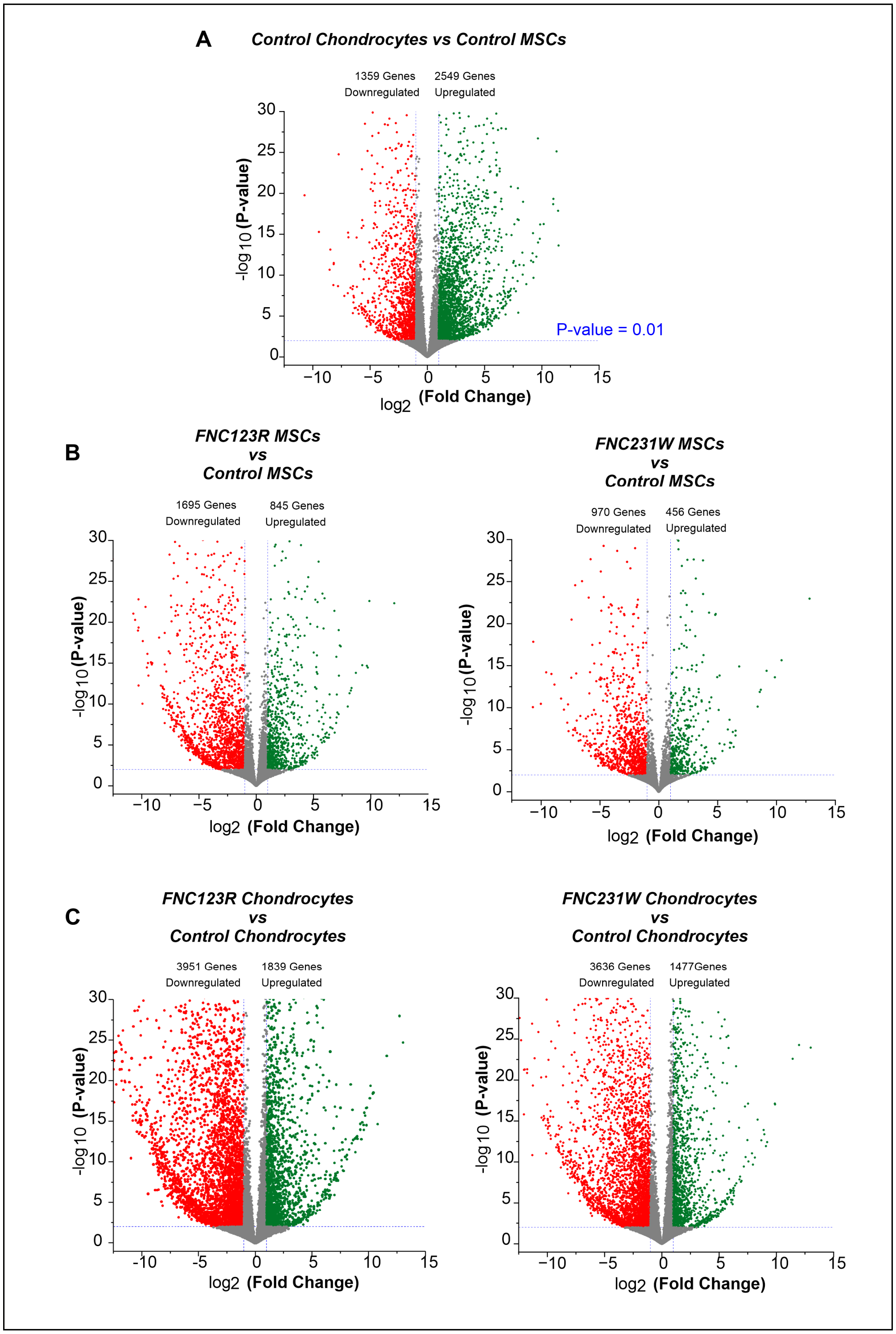


**Supp. Figure 2. FN mutants display a global downregulation of cellular transcriptome.**

**(A)** Volcano plot of the overall differentially expressed genes (DEGs) in control chondrocytes versus control MSCs. Note the higher number of upregulated genes during chondrogenesis. **(B,C)** Volcano plots show the overview of DEGs in FN mutants versus control at the MSC **(B)** and the chondrocyte stage **(C)**.

### Supplemental Figure 3

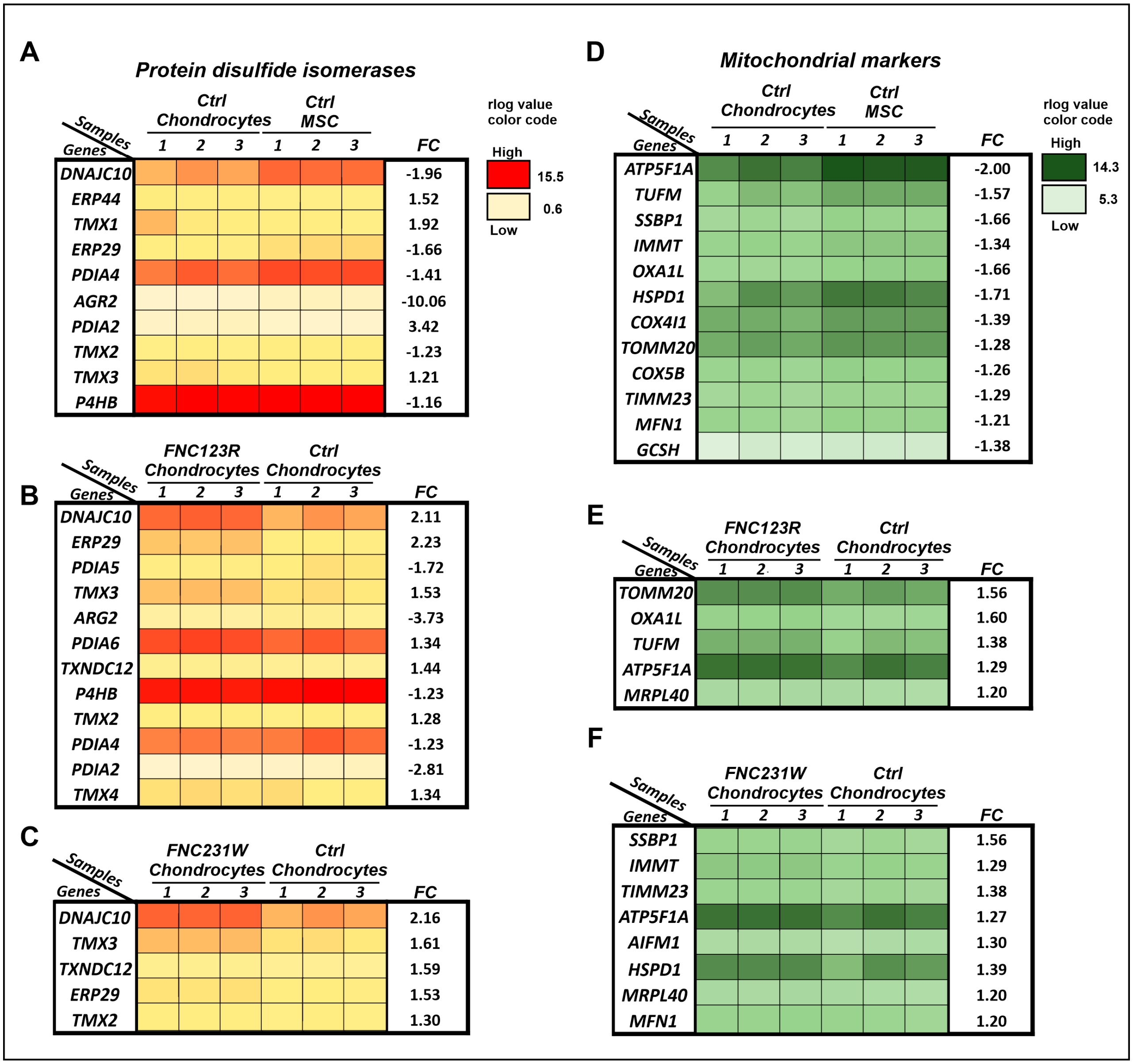


**Supp. Figure 3. SMDCF cells are characterized by misfolded protein response and alterations in mitochondrial markers.**

Heat maps of rlog counts obtained from the RNAseq analysis of differentially expressed markers in the protein disulfide isomerase family **(A-C)**, or of mitochondrial markers **(D-F)**. Data show the differentially expressed genes in control chondrocytes versus control MSCs **(A,D)**, or in FN mutant chondrocytes versus control chondrocytes **(B,C,E,F)**. The data were obtained from 3 technical replicates per sample group. Statistical analysis was performed using the nbinomWaldTest followed by the Benjamini-Hochberg method. For each group, only genes with a significance of $p \leq 0.05$ are shown in descending order, with the lowest p-values on the top. The color codes for the rlog values are indicated on the right. FC represents fold-change.

#### Supplemental Figure 4

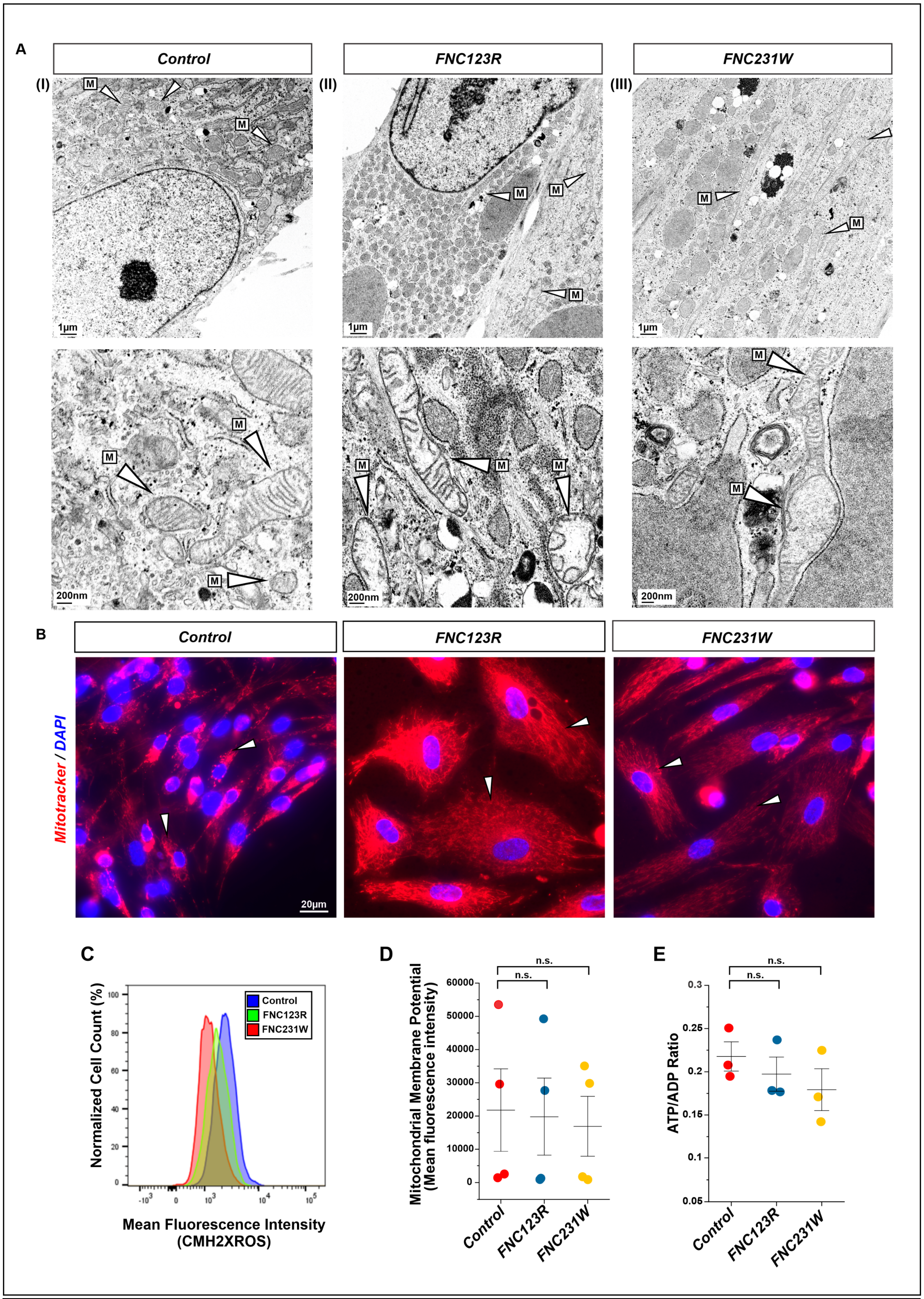


**Supp. Figure 4. Analysis of mitochondrial structure and functional parameters.**

**(A)** TEM images of control and SMDCF mutant MSCs at different magnifications in the top and bottom panels showing mitochondria (M, arrowheads). **(A-I)** Control cells exhibit normal sized mitochondria with typical cisternae structure and distribution. **(A-II)** FN$_{C123R}$ and **(A-III)** FN$_{C231W}$ MSCs display elongated, damaged, and swollen mitochondria with broken cisternae. Scale bars represent 1 µm for the top and 200 nm for the bottom panels. (***continued on next page***)

**(B)** Mitotracker Deep Red FM staining on control and SMDCF mutant MSCs. Mitochondria stained in red (arrowheads) show a typical perinuclear arrangement of mitochondria in control MSCs. Both $FN_{C123R}$ and $FN_{C231W}$ exhibit elongated large mitochondria that are spread across cytoplasm. DAPI (blue) is used as a nuclear counter stain in all images. Scale bar represents 20 µm for all images. **(C)** Flow cytometry analysis of control versus FN mutant MSCs (normalized to mode) after staining with CMH2XROS an indicator for mitochondrial membrane potential. Shown is the mean fluorescence intensity versus total cell count. **(D)** The graph represents quantification of the mean fluorescence intensities shown in **(C)** of four independent experiments (n=4). The error bars represent ± SEM. Significance was evaluated by one-way ANOVA with Bonferroni post-test. **(E)** ATP/ADP ratio analysis in control and FN mutant MSCs. Each data point represents an average of technical triplicates from an independent experiment of a total of three independent experiments (n=3). Error bars represent ± SEM. Significance was evaluated by one-way ANOVA with Bonferroni post-test.

### Supplemental Figure 5

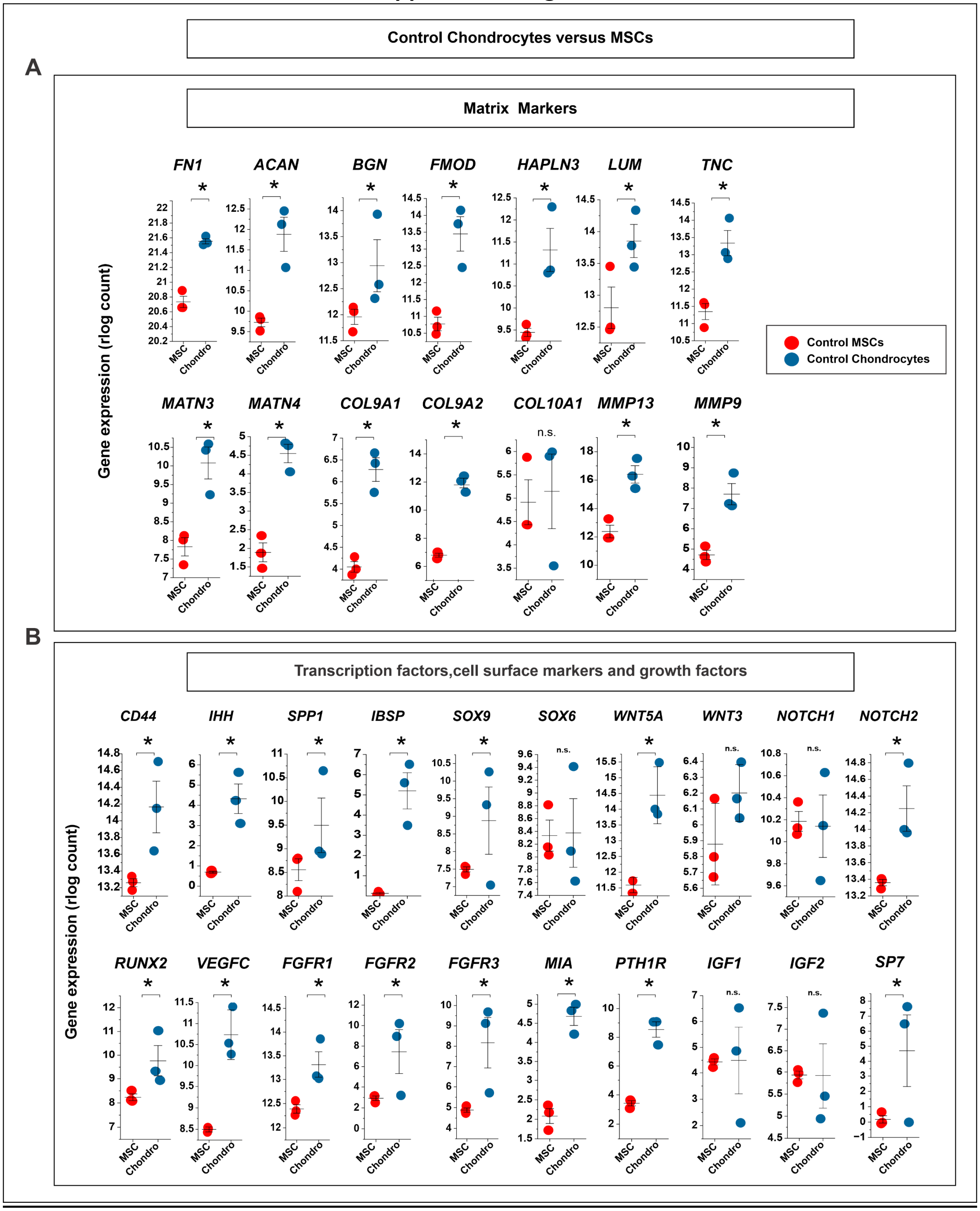


**Supp. Figure 5. Control iPSC-derived stem cells undergo chondrogenesis with increased expression of chondrogenic markers.**

**(A-C)** Graphs represent key chondrogenic markers obtained by RNAseq analysis comparing control chondrocytes versus control MSCs. The transcript reads for each marker are represented as rlog counts and sorted into **(A)** matrix markers, and **(B)** transcription factors, cell surface markers, and growth factors. Data points represent 3 technical replicates per sample group. Statistical analysis was performed using the nbinomWaldTest followed by the Benjamini-Hochberg method. $p \leq 0.05$ was taken as significant for all data sets.

### Supplemental Figure 6

A

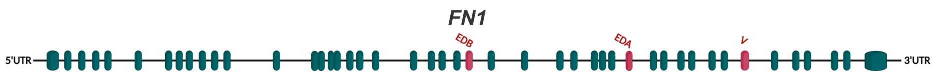


| FN Isoform | Transcript type |
|---|---|
| FN1-201(V+EDA+EDB) | Protein coding |
| FN1-202(V+EDA) | Protein coding |
| FN1-203(V+EDA+EDB) | Protein coding |
| FN1-204(V) | Protein coding |
| FN1-205 | Protein coding |
| FN1-206 (EDA + V) | Protein coding |
| FN1-207(V) | Protein coding |
| FN1-208 | Protein coding |
| FN1-209(EDB) | Protein coding |
| FN1-210(V) | Protein coding |
| FN1-211(V) | Protein coding |
| FN1-212(V+EDA) | Protein coding |
| FN1-213(V) | Protein coding |
| FN1-223 | Non coding |

| FN Isoform | Transcript type |
|---|---|
| FN1-214 | Retained intron |
| FN1-215 | Retained intron |
| FN1-216 | Retained intron |
| FN1-217 | Retained intron |
| FN1-218 | Retained intron |
| FN1-219 | Retained intron |
| FN1-220 | Retained intron |
| FN1-221 | Retained intron |
| FN1-222 | Retained intron |
| FN1-224 | Retained intron |
| FN1-225 | Retained intron |
| FN1-226 | Retained intron |
| FN1-227 | Retained intron |

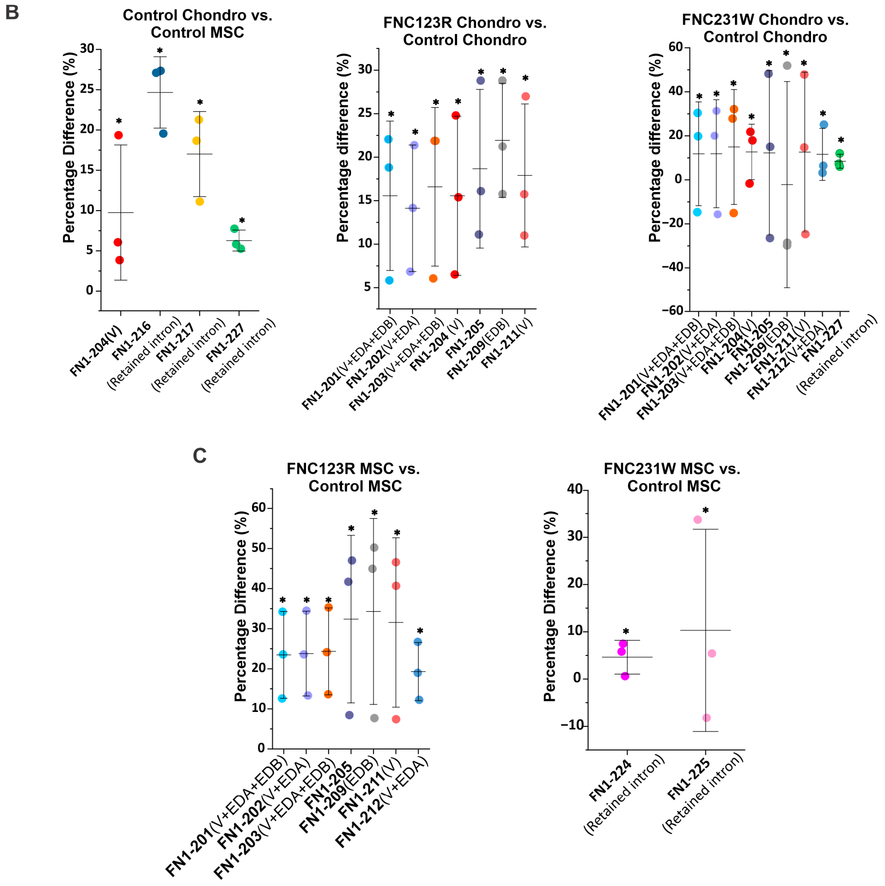


**Supp. Figure 6. FN mutants are characterized by differential expression of FN isoforms during chondrogenesis.**

**(A)** Schematic representation of the exons comprising *FN1* gene transcripts based on the ensemble database (https://www.ensembl.org). Frequently spliced exons are color-labeled in red. The table summarizes the 27 known alternatively spliced FN mRNAs, which were present in all analyzed samples. **(B)** The graphs represent the percentage difference of FN isoform levels that are significantly altered in control chondrocytes versus controls MSCs or in FN mutant chondrocytes versus control chondrocytes. **(C)** Data represent the significantly altered FN isoforms in FN mutant MSCs versus control MSCs. All data points represent 3 technical replicates per sample group per gene. Statistical analysis was performed using the nbinomWaldTest followed by the Benjamini-Hochberg. $p \leq 0.05$ was taken as significant for all data sets.

#### Supplemental Figure 7

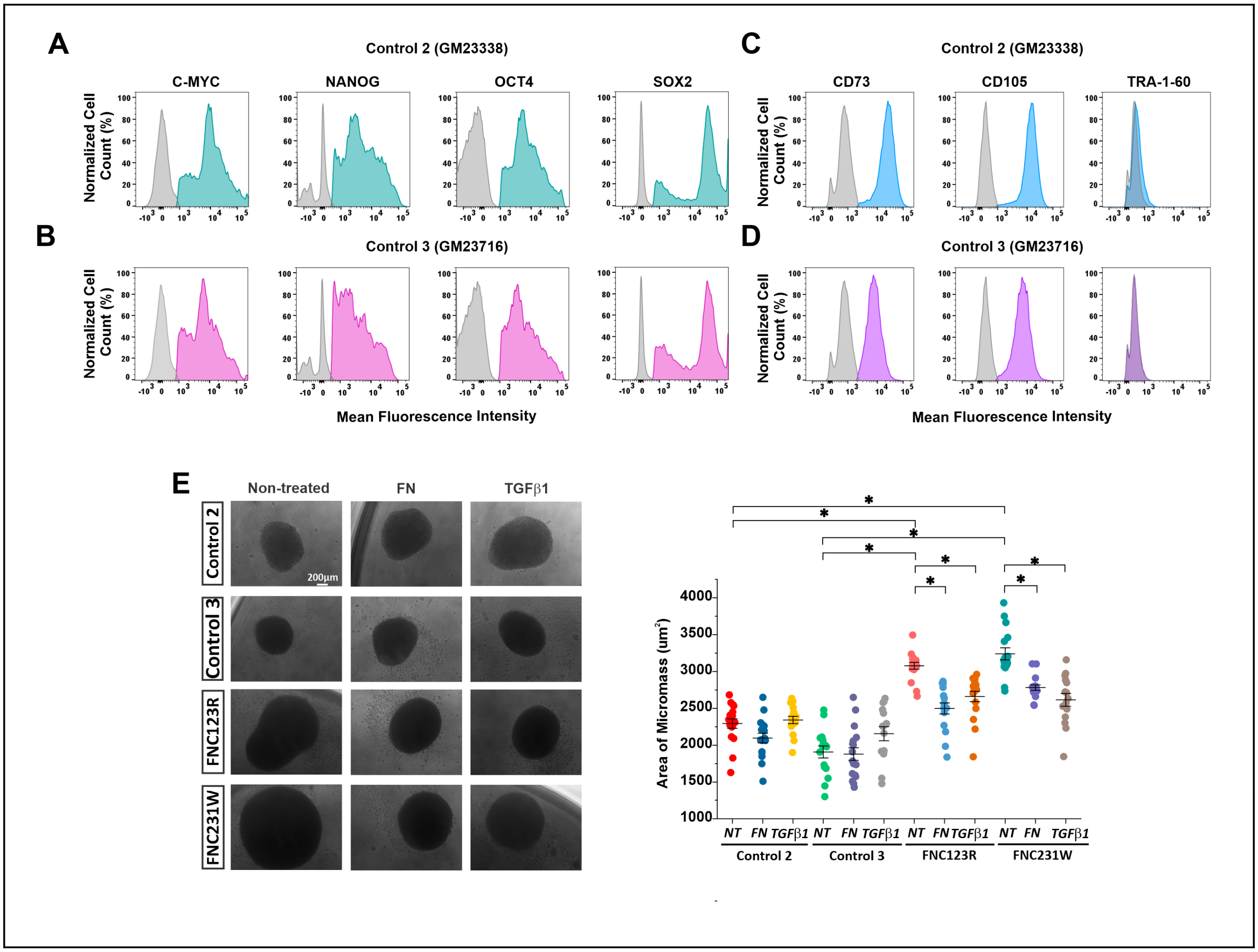


**Supp. Figure 7. Validation of rescue experiment of delayed condensation using additional iPSC control cells.**

The data shown in **Figure 11E,F** were further validated with additional iPSC controls (Ctrl2, GM23338, and Ctrl3, GM23716) obtained from the Coriell Institute for Medical Research. **(A-D)** Base characterization of the additional control iPSCs. **(A,B)** Flow cytometry analyzing the expression of pluripotent markers, C-MYC, NANOG, SOX2, and OCT4. Positive cell populations are represented in green for Ctrl2 iPSCs **(A)** and in magenta for Ctrl3 iPSCs **(B)**. **(C,D)** iPSC-derived differentiated MSC controls were tested by flow cytometry for the presence of MSC markers CD73 and CD105 and the absence of the undifferentiated marker TRA-1-60. Positive cell populations are represented in blue for Ctrl2 MSCs **(C)** and in pink for Ctrl3 MSCs **(D)**. Fluorescence minus one (FMO, gray) for each of the antibodies represents the negative control per cell group in all conditions. **(E)** Representative images of live cell condensation assays at 6 d for the iPSC-derived MSC Ctrl2 and Ctrl3 versus FN mutant cells either non-treated (NT) or treated with FN (25 µg/mL) or TGFβ1 (10 ng/mL). Quantification of micromass areas as shown in **(E)**. The graph represents data from three independent experiments with a total of 15-18 micromasses per sample group. The error bars represent ±SEM. Statistical analysis was performed with the unpaired Student's two-tailed t-test with Welch's correction and test for equal variance (F-test). The scale bar represents 200 µm.

***Supplemental Table 1.*** *List of antibodies used in this study.*

| *List of Antibodies* | *Source* | *Catalogue Number* | *Experimental Use** |
|---|---|---|---|
| Anti-human FN antibody | Millipore Sigma | F3648 | IB (1:1000) and IS (1:500) |
| Anti-human FN antibody (P) | from collaborator | n.a. | IG (1:500) |
| Anti-human FN antibody (M) | from collaborator | n.a. | IG (1:200) |
| Anti-EDA FN antibody | Millipore Sigma | SAB4200784 | IB (1:1000) and IS (1:1000) |
| Anti-GAPDH antibody | Cell Signaling | 2118S | IB (1:1000) |
| Anti-Rab5 antibody | Cell Signaling | 3547T | IS (1:200) |
| Anti-Rab7 antibody | Cell Signaling | 9367T | IS (1:200) |
| Anti-SRP6 antibody | Cell Signaling | 2217T | IS (1:200) |
| Anti-GM103 antibody | BD Biosciences | 610822 | IS (1:200) |
| Anti-HSP47 antibody | Enzo Life Sciences | ADI-SPA-470 | IS (1:500) |
| Anti-LAMP1 antibody | Abcam | ab24170 | IS (1:500) |
| Anti-LC3 antibody | Invitrogen | PA1-16930 | IS (1:200) |
| Anti-GRP78/BIP antibody | Abcam | ab109659 | IS (1:500) |
| Anti-ATG5 antibody | Proteintech | 10181-2-AP | IS (1:200) |
| Anti-Collagen type 2 antibody | DSHB | II-II6B3 | IS (1:20) |
| Anti-human CD73 antibody clone AD2 FITC conjugated | Invitrogen | 17-1057-41 | FC (1:300) |
| Anti-human TRA-1-60 antibody clone TRA-1-60R PE conjugated | StemCell Technologies | 60064PE.1 | FC (1:300) |
| Anti-human/Mouse SOX2 antibody eFluor 660 conjugated | Invitrogen | 50-9811-82 | FC (1:300) |
| Anti-NANOG antibody (23D2-3C6) DyLight488 conjugated | Invitrogen | MA1-017-D488 | FC (1:300) |
| Anti-cMYC antibody Coral lite 555 conjugated | Invitrogen | CL555-10828 | FC (1:300) |
| Anti-OCT4 antibody Coral lite 594 conjugated | Proteintech | CL594-11263 | FC (1:300) |
| Goat anti-Rabbit IgG (H+L) Cross-Adsorbed Secondary Antibody, Alexa Fluor 488 | Invitrogen | A-11008 | IS (1:200) |
| Goat anti-Mouse IgG (H+L) Cross-Adsorbed Secondary Antibody, Cyanine5 | Invitrogen | A10524 | IS (1:200) |
| Goat anti-Mouse IgG (H+L) Cross-Adsorbed Secondary Antibody, Cyanine3 | Invitrogen | A10521 | IS (1:200) |
| Peroxidase-Conjugated AffiniPure Goat Anti-Mouse IgG (H+L) | Jackson ImmunoResearch | 115-035-003 | IB (1:1000) |
| Peroxidase-Conjugated AffiniPure Goat Anti-Rabbit IgG (H+L) | Jackson ImmunoResearch | 111-035-003 | IB (1:1000) |
| 12 nm Colloidal Gold AffiniPure Donkey Anti-Rabbit IgG (H+L) (EM Grade) | Jackson ImmunoResearch | 711-205-152 | IG (1:20) |
| 18 nm Colloidal Gold AffiniPure Goat Anti-Mouse IgG (H+L) (EM Grade) | Jackson ImmunoResearch | 115-215-166 | IG (1:20) |

** IB: Immunoblotting, IS: Immunostaining, IG: Immunogold labelling, FC: Flowcytometryss*

***Supplemental Table 2.*** *Primers specific for FN1 Sanger sequencing and qPCR analysis.*

| Annealing position (nt)* | ***Orientation*** | ***Sequence 5'-3'*** | ***Experimental Use*** |
|---|---|---|---|
| 198 | Sense | GCGGACCTACCTAGGCAATG | Validation of c.367T>C |
| 569 | Antisense | CACATAGGAAGTCCCAGCAG | |
| 264 | Sense | GAGTAAACCTGAAGCTGAAGAG | Validation of c.693C>G |
| 931 | Antisense | GAGTAGACCACACCACTGTC | |
| s5162 | Sense | CCAACATTGATCGCCCTAAAG | qPCR |
| 5275 | Antisense | GATTCCATCCTCAGGGCTC | qPCR |

* Numbers refer to the nucleotide (nt) position in the FN1 sequence (NM_212482.4) that anneals with the first 5' nt of the primer.
